# SKiM - A generalized literature-based discovery system for uncovering novel biomedical knowledge from PubMed

**DOI:** 10.1101/2020.10.16.343012

**Authors:** Kalpana Raja, John Steill, Ian Ross, Lam C Tsoi, Finn Kuusisto, Zijian Ni, Miron Livny, James Thomson, Ron Stewart

## Abstract

Literature-based discovery (LBD) uncovers undiscovered public knowledge by linking terms A to C via a B intermediate. Existing LBD systems are limited to process certain A, B, and C terms, and many are not maintained. We present SKiM (Serial KinderMiner), a generalized LBD system for processing any combination of A, Bs, and Cs. We evaluate SKiM via the rediscovery of discoveries by Don Swanson, who pioneered LBD. Using only literature from the 19th century up to a year before Swanson’s discoveries, SKiM uncovers all five discoveries. We apply SKiM to repurposing drugs for 26 conditions of high prevalence. Manual analysis confirmed 65 discoveries useful for four diseases from Swanson’s discoveries from one to 31 years prior to their first validation by clinical trials. SKiM predicts many new potential drug candidates representing prime targets for wet lab validation. SKiM can be applied to any biomedical inquiry sufficiently mentioned in the literature.

## Introduction

Thousands of articles are added to PubMed daily with the total number of articles approaching 31 million (M)^1^. It is challenging for researchers to keep up-to-date with all relevant information within their domain of interest, and it is very difficult to keep track of discoveries made outside their domain. Furthermore, uncovering hidden knowledge by associating topics across domains is extremely difficult for humans, except on a very limited scale.

Biomedical text mining applied to all ~31M PubMed abstracts can help expedite new scientific discoveries^2,3^. One text mining strategy is literature-based discovery (LBD), which uncovers undiscovered public knowledge by associating topics across domains^4^. Swanson pioneered LBD by combining two known pieces of information to derive a novel connection. He hypothesized dietary fish oil as a remedy for Raynaud’s disease (RD) from two sets of PubMed abstracts, one set that identifies high blood viscosity as a condition in RD, and the other set that suggests that dietary fish oil can be useful to lower blood viscosity^5^. Swanson’s discovery was later validated through clinical trials^6^. Automated LBD has become a core text mining task^7^. The form of LBD we implement here is the “open discovery A➔B➔C model,” which takes only a concept of interest (the “A” term) and retrieves a list of associated concepts (“C” terms) through some intermediate concepts (“B” terms). The application of LBD to the scientific community is limited due to the lack of influential methodology^9,10^. Existing LBD systems fall into one of the following categories: co-occurrence models, semantic models, graph-based models, distributional models, and hybrid models^7,11^. While many LBD systems have been built^7,8,10,12^, only BITOLA^8^ and LION LBD^10^ have a functioning web interface. BITOLA^8^ is restricted to use the semantic types from the Unified Medical Language System (UMLS) Metathesaurus^13^ as B and C terms. Drug repurposing with BITOLA is incomplete, as the available controlled vocabularies do not contain a comprehensive list of drugs and do not include many drugs and synonyms from DrugBank^14^ and PharmGKB^15,16^. LION LBD is specific to cancer biology, and restricted to use only chemicals, diseases, mutations, genes, cancer hallmarks, and species as B and C terms^10^. The system is not flexible to run on specific types of concepts as B (e.g. phenotypes or genes) and C terms (e.g. drugs). For this reason, LION LBD is not applicable for drug repurposing. Many earlier LBD systems queried the Medical Subject Heading (MeSH) index because it is a structured text and the vocabularies are controlled^8,17^. The approach is not effective for two major reasons: *(1)* only ~24M PubMed abstracts are indexed with MeSH, and *(2)* all significant subject terminologies from the abstracts may not be present in MeSH^18^. A pattern-based approach extracted disease-gene and gene-drug relationships from PubMed abstracts for repurposing drugs for different cancers^4^. The patterns generated are domain specific and may not perform well on other domains.

We present here SKiM (**S**erial **Ki**nder**M**iner), an open discovery, domain-independent LBD system for processing any combination of A, B, and C terms from PubMed titles and abstracts. SKiM overcomes the limitations of the existing LBD systems previously described. SKiM builds upon the features of KinderMiner^19^, our recently developed text mining system to filter statistically significant co-occurring concept pairs from PubMed abstracts. Many existing LBD systems^10,20,21^ and popular text mining systems such as PolySearch2^2^ use a co-occurrence approach for extracting information from PubMed. KinderMiner gave interesting and promising results on identifying transcription factors for cell reprogramming, retrieved potential drugs for lowering blood glucose^19^, performed better than STRING^22^ and PolySearch2^2^ on retrieving mitochondrial protein-protein interactions (PPI)^23^, identified lab tests that can be repurposed for other diagnostic goals^24^, and identified phenotypes associated with FMR1 premutation^25^. In the current study, we applied SKiM on repurposing drugs for four diseases from the research articles published by Swanson and his colleagues^5,26–29^: RD, migraine, Alzheimer’s disease (AD), and schizophrenia. Drug repurposing is a universally agreed-upon need to maximize resources and quickly respond to new challenges^30–33^.

Importantly, we compiled two comprehensive lexicons: *(1)* a phenotypes and symptoms lexicon and *(2)* a drugs lexicon (see Methods for details). The lexicons are critical to allowing SKiM to have as wide a view as possible of potential A to B to C connections. We further applied SKiM to 22 additional high-impact conditions. SKiM identified candidate drugs up to 31 years prior to the first validation by clinical trials published in PubMed, and up to 32 years prior to approval by the US Food and Drug Administration (FDA)^34^. We provide instructions for building a local text index and software so that users can run SKiM locally.

## Results

### Overview

For a given concept A, SKiM uncovers associated C terms through intermediate B terms (A➔B➔C) (Fig. 1). SKiM counts PubMed titles and abstracts with the co-occurring A-B or B-C pairs and filters statistically significant pairs using the one-sided Fisher Exact Test (FET) p-value. Unless otherwise stated, a “discovery” or “finding” is defined as targets that achieve an FET p-value of less than 1×10^-05^. We evaluated SKiM on rediscovering the five discoveries by Swanson and his colleagues by allowing SKiM to only “read” papers up to one year prior to Swanson’s discoveries. We further applied SKiM for repurposing drugs for four diseases studied by Swanson as well as 22 additional high-impact conditions. We used our comprehensive phenotypes and symptoms list as the B terms, and a subset of drugs annotated with diseases, genes, phenotypes, variants, or haplotypes from various expert curated resources as C terms. We evaluated the repurposed drugs on our expert curated disease-drug associations (DDA). By manual analysis of PubMed articles, clinical trials databases, and FDA databases, we showed discoveries of many drugs useful for the diseases up to several years prior to validation.

**Fig 1.**
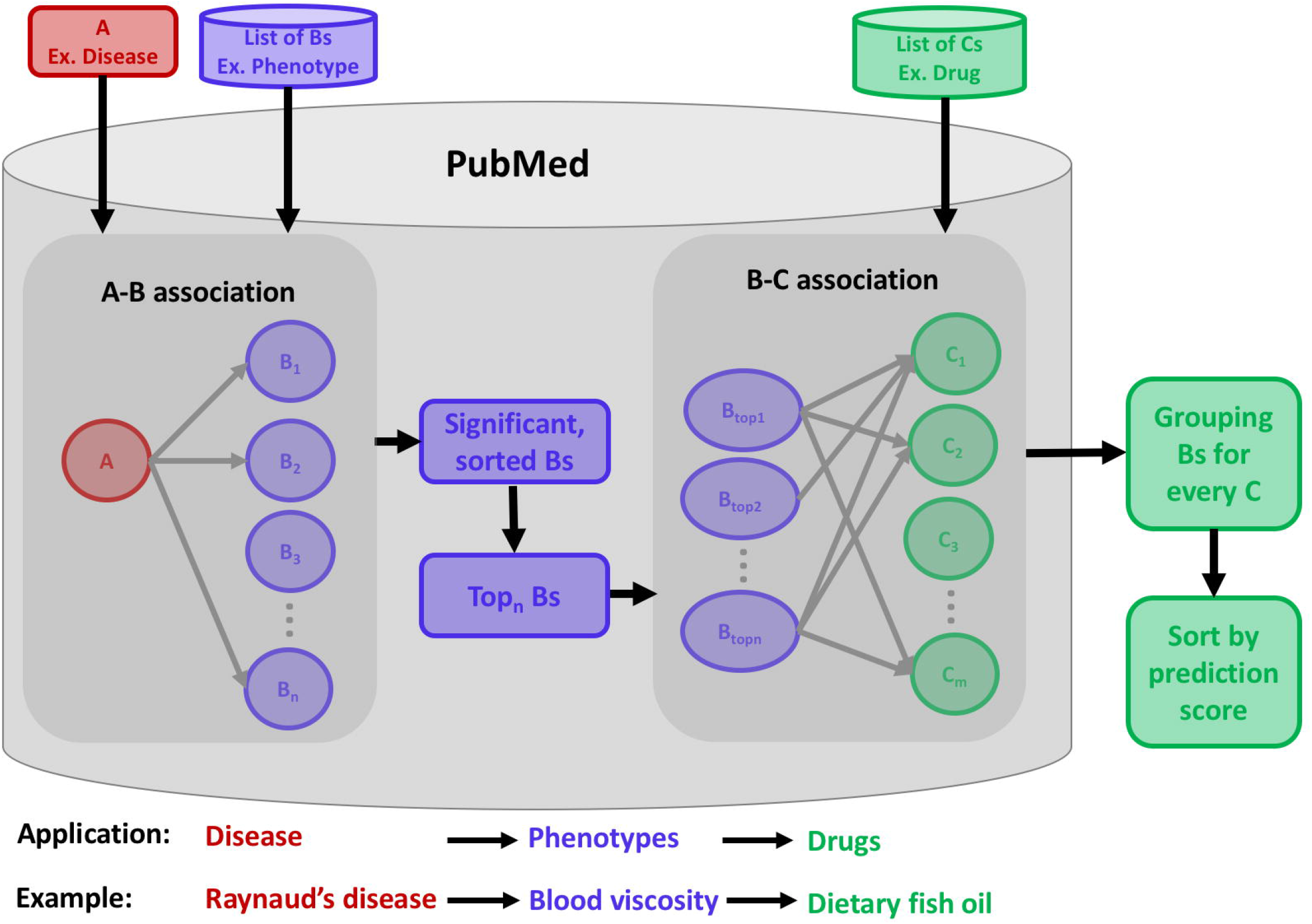
SKiM Workflow. SKiM executes in two levels. First, it finds significant B terms associated with the input A term (see A-B association). Second, it finds significant C terms for each top n B terms from A-B association (see B-C association). B-C findings are grouped together and sorted based on the prediction score. For drugs repurposing application, A term is a disease of interest, B terms are a comprehensive list of phenotypes and symptoms, and C terms are drugs.

### Evaluation based on Swanson’s five prior discoveries

When SKiM was allowed to read PubMed abstracts published one year prior to the discovery, SKiM uncovered all the discoveries at a FET p-value less than 1×10^-05^ (the default p-value used by KinderMiner previously^19^) (Fig. 2A, Supplementary Data 1). Simulation indicates that a p-value cut-off of 1×10^-05^ is conservative enough to limit the false discovery rate (FDR) for A to C to below 0.005 while still maintaining a power > 0.7. Given that SKiM is primarily designed to find targets for further study, having some false negatives is a reasonable tradeoff for a low FDR. We don’t perform ordinary FDR correction as the distributions of the p-values violate the model assumptions of most state-of-the-art FDR control methods due to the unbalance between small abstract hit counts and enormous total number of PubMed abstracts^35^ (see Supplementary Method 1). Swanson and his colleagues identified drug, hormone, inflammation, and other concepts as B terms for uncovering the discoveries^26^. SKiM rediscovered all the discoveries by using phenotypes and symptoms as B terms (Supplementary Note 1) and many novel discoveries. The results show that the phenotypes and symptoms as B terms are appropriate for drug repurposing and will likely be useful for other tasks.

**Fig 2.**
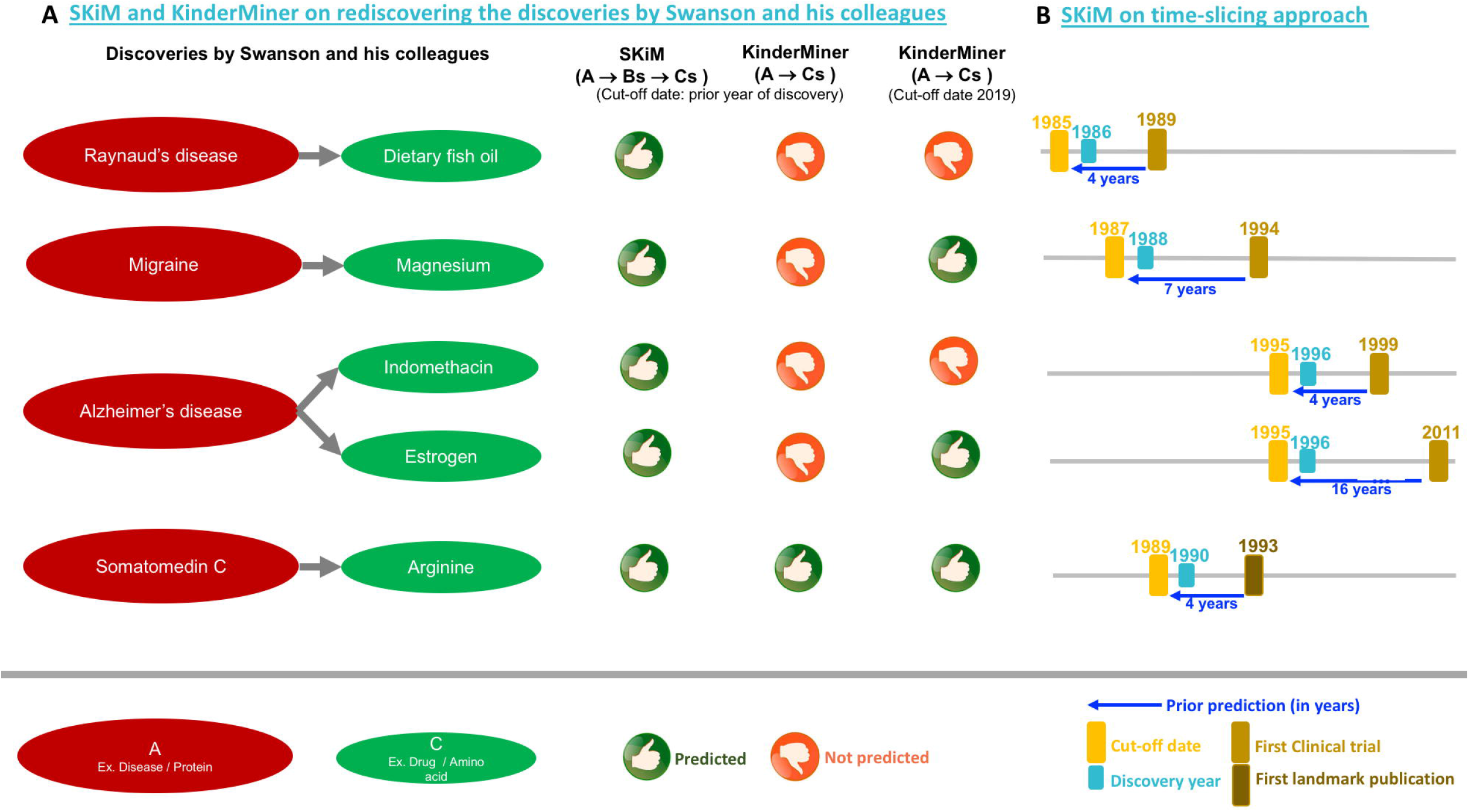
Performance on the discoveries by Swanson and his colleagues. **A.** SKiM and KinderMiner on rediscovering the discoveries by Swanson and his colleagues. SKiM rediscovered all five discoveries at the cut-off date one year prior to the discoveries by Swanson and his colleagues. KinderMiner rediscovered only one discovery at the cut-off date one year prior to the discoveries by Swanson and his colleagues. KinderMiner failed to rediscover two discoveries when the cut-off date is relaxed to March 2019. **B.** Performance of SKiM on time-slicing approach. SKiM uncovered five discoveries by Swanson and his colleagues up to 16 years prior to validation by clinical trials and the first landmark publication.

Discoveries by Swanson and his colleagues were later validated by clinical trials or published as a new discovery (Supplementary Data 1). By uncovering all the discoveries by Swanson and his colleagues, SKiM demonstrates the ability to uncover the discoveries from PubMed abstracts prior to the validation by clinical trials or the first landmark publication (Fig. 2B).

### Comparison with KinderMiner and existing LBD systems

First, we compared SKiM with KinderMiner on five discoveries by Swanson and his colleagues. When KinderMiner was allowed to read PubMed abstracts published one year prior to the discovery, KinderMiner (which looks for direct A➔C associations) extracted only one discovery at the default FET p-value (Fig. 2A, Supplementary Data 1). KinderMiner did not improve much when the FET p-value is relaxed or cut-off date is extended to include abstracts past the date of initial discovery (Fig. 2A, Supplementary Table 1). Thus, SKiM (A➔B➔C) performs better than KinderMiner (A➔C) with regard to uncovering Swanson’s discoveries.

Next, we compared SKiM with the existing LBD systems on five discoveries by Swanson and his colleagues. Existing LBD systems are restricted to process certain combinations of A, B, and C terms^10,35^. BITOLA is restricted to use only UMLS concepts as A, B, and C terms^8^. LION LBD is restricted to use only chemicals, diseases, mutations, genes, cancer hallmarks, and species as A, B, and C terms^10^. We tested the open discovery model of BITOLA and LION LBD on rediscovering five discoveries by Swanson and his colleagues. BITOLA rediscovered only three of five discoveries by Swanson and his colleagues at an unknown cut-off date (and thus may be unfairly benefiting from papers published after Swanson’s discoveries) (Supplementary Note 2). LION LBD uncovered none of the C terms from the discoveries by Swanson and his colleagues at the cut-off date of one year prior to the discovery (Supplementary Note 3). Moreover, LION LBD is specific to the case of molecular biology of cancer^10^ and thus is not reliable for drug repurposing because there is no option to select specific type of B terms (e.g. genes or phenotypes) and C terms (e.g. drugs).

### Application of SKiM to drug repurposing

In the prior section, we focused on applying SKiM to attempt to replicate Swanson’s five discoveries using a cut-off date of one year before Swanson’s discoveries and focusing on just the replication of Swanson’s five discoveries. Here, we further allow SKiM to search for additional candidates for drug repurposing for four diseases from Swanson’s work, RD, migraine, AD, and schizophrenia using both a cut-off date of year before Swanson’s discoveries as before, and also relaxing the cut-off date to allow SKiM to “read” abstracts through March 2019. We used 9,272 phenotypes and symptoms as B, 9,665 drugs as C, and considered the top 50 Bs from A➔B for uncovering B➔C. Considering all Bs from A➔B for uncovering B➔Cs is computationally expensive. A common practice is to consider only some number of the top ranking Bs from A➔B^8^, and we follow that practice here.

We evaluated SKiM findings for RD, migraine, AD, and schizophrenia on our expert curated DDA compiled from four existing resources (Supplementary Table 2, see Methods for details). In addition, we manually analyzed SKiM findings and compared them with real-world findings in four ways. First, we assessed how many years in advance SKiM found drug-disease pairs that were later mentioned together in a PubMed abstract. Second, for drugs which became a published treatment use, we assessed how many years in advance of that publication SKiM found that drug. Third, we assessed how far in advance SKiM found a pair that later appeared in a clinical trial. Fourth, we assessed how far in advance of FDA approval SKiM was able to identify drug-disease pairs. We describe each of these methods in greater detail in the following sections.

### Evaluation using expert curated disease-drug associations

SKiM uncovered many drugs from our expert curated DDA (Supplementary Data 2). The recall achieved by SKiM is higher than the recall achieved by KinderMiner on all four diseases (Fig. 3A, Supplementary Note 4). In text mining, precision at top N predictions, where N is the count of included predictions (i.e. precision@N) is a common approach to evaluate systems such as SKiM and KinderMiner^36^. The precision achieved by SKiM at top 20 drugs (i.e. precision@20) (Supplementary Data 3) is higher than the precision@20 achieved by KinderMiner (Fig. 3B, Table 1, Supplementary Note 5). Interestingly, SKiM uncovered many drugs only through A➔B and B➔C (Table 2). These drugs are missing in the DDA and these new findings are interesting ones for further study. Our results suggest the importance of automated approaches like SKiM to uncover unknown public knowledge from PubMed.

**Fig 3.**
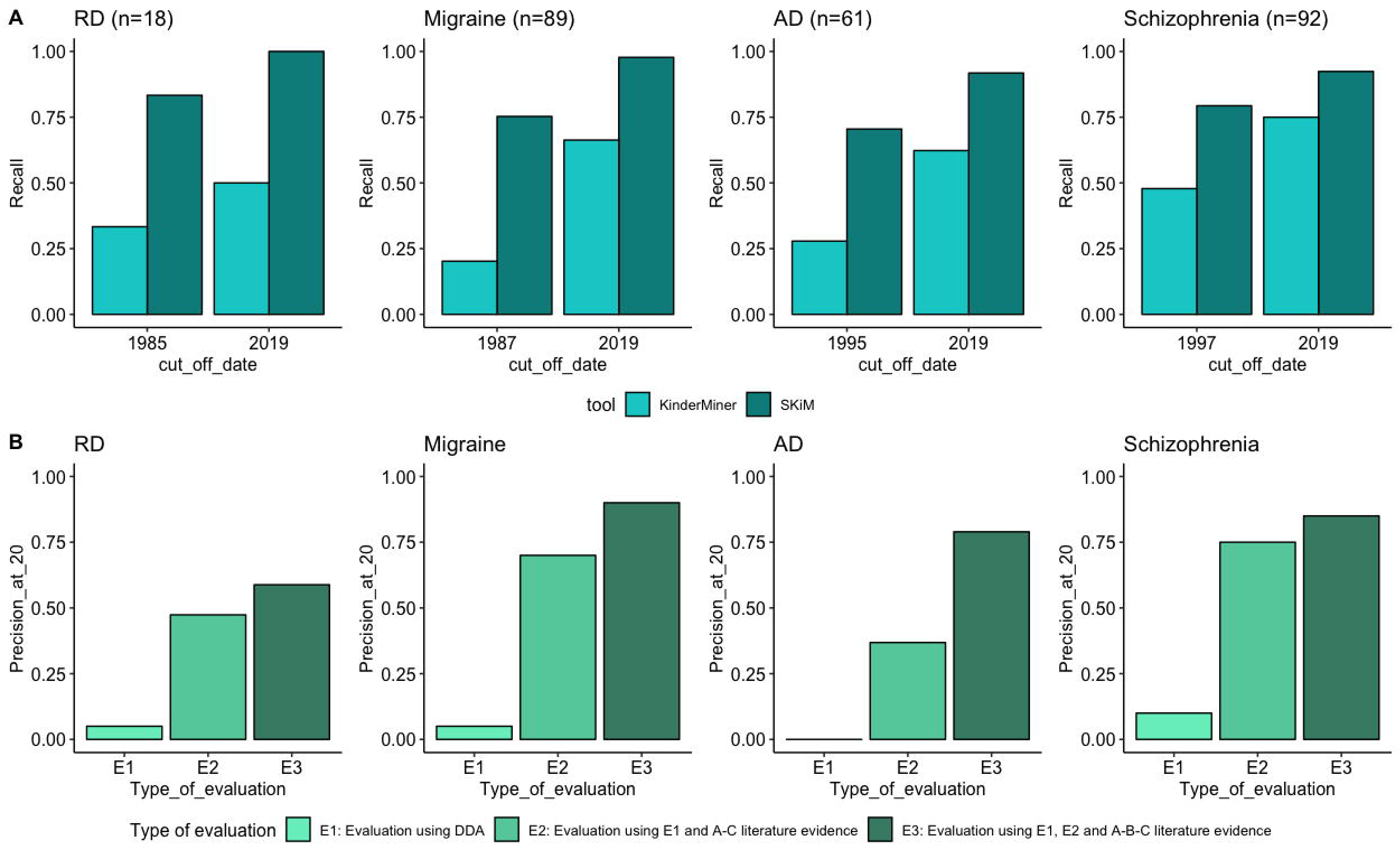
Performance on expert curated DDAs. **A.** Recall achieved by KinderMiner and SKiM on the expert curated DDAs at the cut-off date one year prior to the discoveries by Swanson and his colleagues, and the cut-off date March 2019. There are 18 DDAs for RD, 89 DDAs for migraine, 61 DDAs for AD, and 92 DDAs for schizophrenia. **B.** Precision achieved by SKiM on top 20 repurposed drugs for RD, migraine, AD, and schizophrenia at the cut-off date one year prior to the discoveries by Swanson and his colleagues. The reported precision is based on the evaluation on the expert curated DDAs and manual curation.

**Table 1.** Performance of SKiM and KinderMiner on the DDA

| Disease | Cut-off date | DDA | KinderMiner |  |  |  |  |  | SKiM |  |  |  |  |  |
| --- | --- | --- | --- | --- | --- | --- | --- | --- | --- | --- | --- | --- | --- | --- |
|  |  |  | TP | FP | FN | R | P | P@20 | TP | FP | FN | R | P | P@20 |
| RD | 1985 | 18 | 6 | 31 | 12 | **0.3333** | 0.1935 | **0.5789** | 15 | 1,604 | 3 | **0.8333** | 0.0094 | **0.5882** |
| migraine | 1987 | 89 | 18 | 46 | 71 | **0.2022** | 0.3913 | **0.7000** | 67 | 1,469 | 22 | **0.7528** | 0.0463 | **0.9000** |
| AD | 1995 | 61 | 17 | 270 | 44 | **0.2787** | 0.0630 | **0.5500** | 43 | 2,667 | 18 | **0.7049** | 0.0165 | **0.7895** |
| schizophrenia | 1997 | 92 | 44 | 78 | 48 | **0.4783** | 0.5641 | **0.5789** | 73 | 1,703 | 19 | **0.7935** | 0.0429 | **0.8500** |

Notes: TP – True positives; FP – False positives; FN – False negatives; R – Recall; P – Precision; P@20 – Precision at top 20 predictions.

**Table 2.**
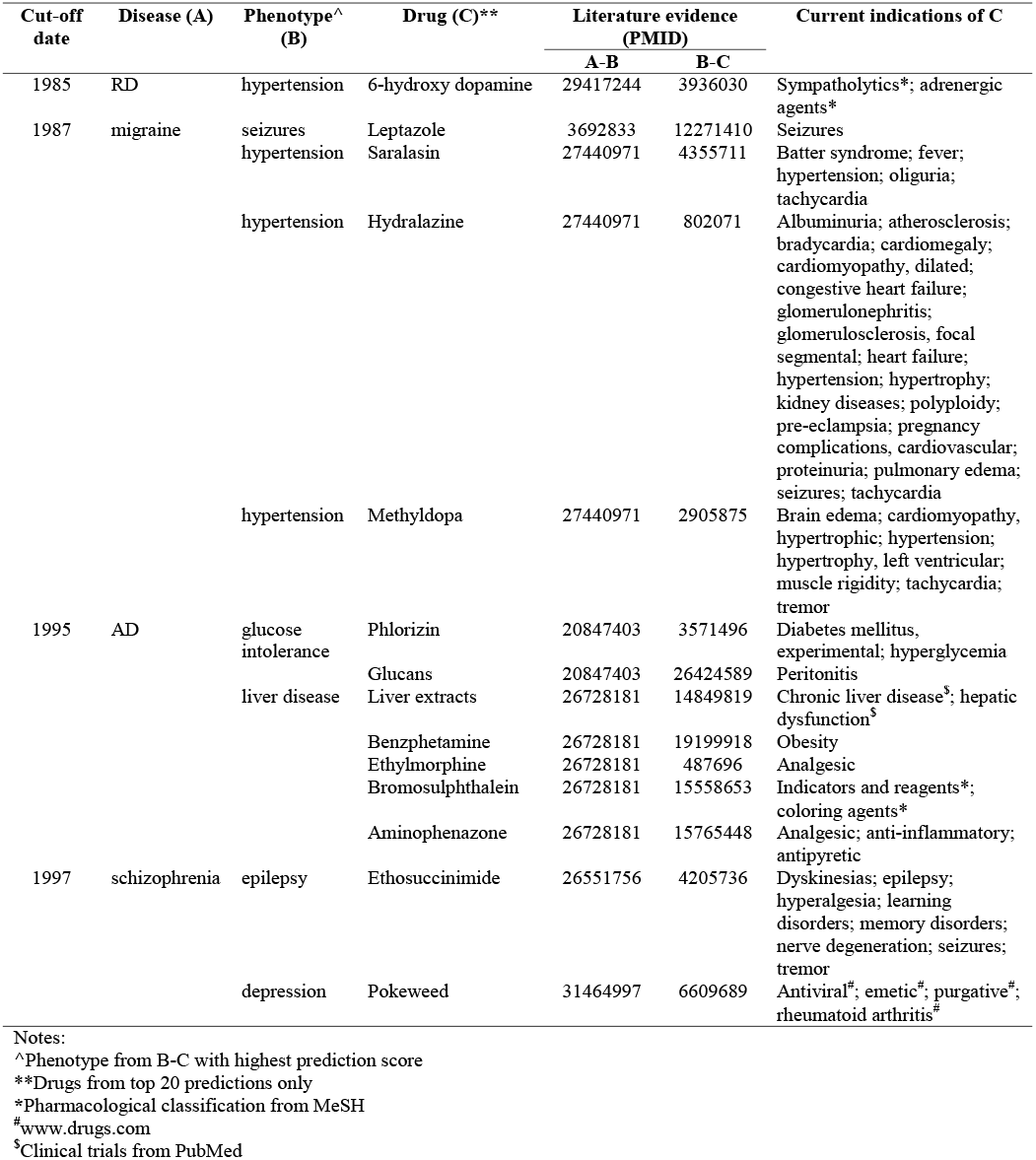
Drugs uncovered only through A→B→C association

Some of the annotated drugs from the DDA for RD, migraine, AD, and schizophrenia are suggested for two or three diseases under study (Supplementary Figure 1). SKiM uncovered most of these drugs for every annotated disease and achieved recall of 0.9333 when it is allowed to read PubMed abstracts until one year prior to the discovery by Swanson and his colleagues (Supplementary Data 4, Supplementary Note 6). Interestingly, SKiM achieved recall of 1.0 when it is allowed to read PubMed abstracts through March 2019 (Supplementary Data 4). The results show that SKiM is sensitive to uncover the right drug for a disease, including cases where a drug is suggested for more than one disease. SKiM does not find all the annotated drugs from the DDA for RD, migraine, AD, and schizophrenia (Supplementary Data 5). This is typically caused by having too few abstracts co-mentioning the A and B terms or B and C terms.

### Evaluation of repurposed drugs at specific cut-off date

Manual analysis on SKiM findings using a particular cut-off date, 1985 for RD, 1987 for migraine, 1995, for AD and 1997 for schizophrenia, shows prior prediction of a drug for a disease. We considered the top 20 SKiM findings for manual analysis and showed promising results via four types of manual analyses as discussed in the following four sections.

### Predictions prior to first co-occurrence in a PubMed abstract

One early indication of an association between a drug and disease is the first co-occurrence in a PubMed abstract. It is important to note that the first co-occurrence in a PubMed abstract does not necessarily indicate the drug is useful for treating the disease and thus is a loose measure of association of drug and disease. A more strict indication of utility may actually come later (based on publications indicating therapeutic use, clinical trial information, or FDA approval as discussed later). If SKiM can predict an association even before this first co-occurrence in a PubMed abstract, this is a strong indication of SKiM’s sensitivity and potential usefulness. For every drug uncovered by SKiM (e.g. Adofen for RD), we manually compared the cut-off date (e.g. 1985) and the year of first co-occurrence in a PubMed abstract (e.g. 1993) to know the number of years (1993 – 1985 = 8 years) SKiM can find a discovery before even the first cooccurrence of the drug and the disease in a PubMed abstract. Intriguingly, SKiM uncovered four drugs for RD, eight drugs for migraine, six drugs for AD, and two drugs for schizophrenia up to 32 years prior to their first co-occurrence with the respective disease in a PubMed abstract (Table 3, Supplementary Data 3). The results show that a wealth of information is available in PubMed and that there is a strong need of automated systems like SKiM for uncovering hidden information even before the first co-occurrence of A and C terms in a PubMed abstract.

**Table 3.** Drugs predicted prior to first publication and first literature suggestion in PubMed

| Cut-off date | Disease (A) | First co-occurrence of A and C in PubMed |  |  | First literature suggesting C for A |  |  |
| --- | --- | --- | --- | --- | --- | --- | --- |
|  |  | Drug (C) | PMID (Year)<br>(Y1) | Prior prediction<br>(Y1 - Cut-off date) | Drug (C) | PMID (Year)<br>(Y2) | Prior prediction<br>(Y2 - Cut-off date) |
| 1985 | RD | Tetrodotoxin | 28089920 (2017) | 32 | Adofen | 11561116 (2001) | 16 |
|  |  | Bupivacaine | 9952101 (1999) | 14 |  |  |  |
|  |  | Adofen | 8434812 (1993) | 8 |  |  |  |
| 1987 | migraine | Methyldopa | 26712075 (2015) | 28 | Methergoline | 9225282 (1997) | 10 |
|  |  | Desoxyphenobarbital | 15246950 (2004) | 17 | ACE inhibitors | 7591740 (1995) | 8 |
|  |  | Leptazole | 10928304 (2000) | 13 | Anticonvulsive agents | 8409107 (1993) | 6 |
|  |  | Enalapril | 8384953 (1993) | 6 | Sodium valproate | 1576648 (1992) | 5 |
|  |  | Quipazine | 1731051 (1992) | 5 | Ergometrine | 2759844 (1989) | 2 |
|  |  | Methergoline | 1330643 (1992) | 5 |  |  |  |
|  |  | Saralasin | 2163854 (1990) | 3 |  |  |  |
|  |  | Labetalol | 3278028 (1988) | 1 |  |  |  |
| 1995 | AD | Aminophenazone | 31797568 (2019) | 24 | Metformin | 21277873 (2011) | 16 |
|  |  | Phlorizin | 26706979 (2015) | 20 | Cod liver oil | 12873849 (2003) | 8 |
|  |  | Benzphetamine | 23904614 (2013) | 18 |  |  |  |
|  |  | Metformin | 16330295 (2005) | 10 |  |  |  |
|  |  | Cod liver oil | 15681801 (2004) | 9 |  |  |  |
|  |  | Ethylmorphine | 12877357 (2003) | 8 |  |  |  |
| 1997 | schizophrenia | (1-34)-human parathyroid hormone | 16526182 (2005) | 8 | Antiepileptic agents | 22466630 (2007) | 10 |
|  |  | 25 hydroxycholecalciferol | 11005548 (2000) | 3 | (1-34)-human parathyroid hormone | 16526182 (2005) | 8 |
|  |  |  |  |  | 25 hydroxycholecalciferol | 17156062 (2003) | 6 |
|  |  |  |  |  | Vigabatrin | 12547302 (2002) | 5 |
|  |  |  |  |  | Insulin-like growth factor I | 11078035 (2000) | 3 |

### Predictions prior to first publication indicating a therapeutic use of a drug for a disease

Here, we evaluated SKiM to determine if it can predict drugs for a disease before the earliest PubMed abstract that indicates the therapeutic use of a drug for treating a disease. Similar to the analysis on first co-occurrence in a PubMed abstract, we restricted our study to the top 20 findings for RD, migraine, AD, and schizophrenia at a cut-off date one year prior to the discoveries by Swanson and his colleagues. For every drug uncovered by SKiM (e.g. Adofen for RD), we manually compared the cut-off date (e.g. 1985) and the year of the first publication indicating the therapeutic use. SKiM can predict a drug for a disease up to 16 years prior to their first publication suggesting the therapeutic use in PubMed (Table 3, Supplementary Data 3).

### Predictions prior to validation by clinical trials

Many discoveries by Swanson and his colleagues were later validated by clinical trials (see Supplementary Note 7 for details). This motivated us to explore SKiM on uncovering the drugs from DDA (positive controls) and new drugs (from top 20 predictions only) prior to their validation by clinical trials in PubMed. SKiM does indeed provide predictions well before clinical trial support. Manual analysis revealed the prediction of 65 drugs (57 drugs from DDA from one to 31 years, and eight new drugs from one to 22 years) prior to the validation by clinical trials (Table 4, Supplementary Note 8). Many drugs from DDA (Supplementary Data 2) and new drugs from top 20 predictions (Supplementary Data 3) were validated by clinical trials at or before the cut-off date used by SKiM. This is expected because the cut-off date is based on the particular discoveries by Swanson and his colleagues and not necessarily the drug from DDA or new drug of interest. Many drugs from DDA and new drugs are not yet validated by clinical trials (Table 5). The drugs listed in Table 5 are candidate drugs for further analysis by wet lab researchers.

**Table 4.**
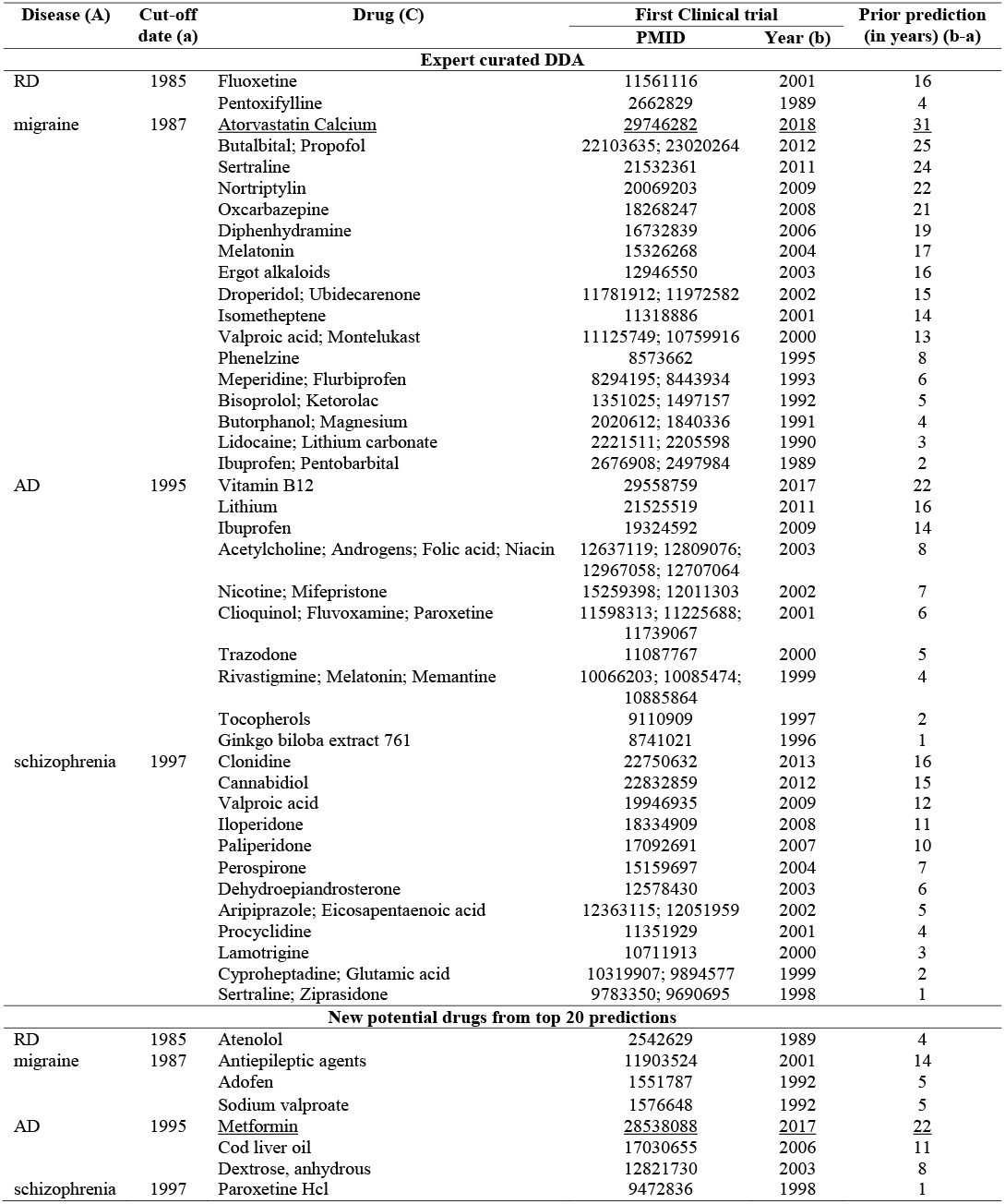
SKiM predictions prior to validation by clinical trials

**Table 5.** SKiM predictions for future validation by clinical trials

| Cut-off date | Condition | DDA | New potential drugs from top 20 predictions |
| --- | --- | --- | --- |
| 1985 | RD | Indomethacin<br>Glucocorticoids<br>Papaverine<br>Tolazoline | 1 Sar 8 Ala Angiotensin II<br>Tetrodotoxin<br>Hydralazine<br>Bupivacaine<br>Phentolamine<br>6-Hydroxydopamine<sup>#</sup> |
| 1987 | migraine | Enzyme inhibitors<br>Phenytoin<br>Naloxone<br>Phenobarbital<br>Imipramine<br>Diflunisal<br>Nalbuphine<br>Thiethylperazine<br>Paroxetine<br>Sulindac<br>Methsuximide<br>Niacin<br>Nicergoline<br>Meclofenamate sodium<br>Abobotulinumtoxina<br>Pergolide | Leptazole*<sup>#</sup><br>Labetalol<br>Enalapril<br>ACE inhibitors<br>Quipazine<br>Saralasin<sup>#</sup><br>Methergoline<br>Hydralazine<sup>#</sup><br>Anticonvulsive agents<br>Desoxyphenobarbital<br>Methyldopa<sup>#</sup><br>Ergometrine |
| 1995 | AD | Dipyridamole<br>Maprotiline<br>Midazolam<br>Amphotericin b<br>Reserpine<br>Ergoloid mesylates<br>Perphenazine<br>Pindolol<br>Goserelin<br>Curcumin<br>Molindone<br>Sulindac sulfide<br>Minocycline | Liver Extracts<sup>#</sup><br>Glucose-6-phosphate*<br>Isotonic NaCl<br>Deoxyglucose*<br>Benzphetamine<sup>#</sup><br>Phlorizin<sup>#</sup><br>5,6-benzoflavone<br>Ethylmorphine<br>Bromosulphthalein*<br>Glucans<sup>#</sup><br>Bromsulphalein*<br>Aminophenazone*<sup>#</sup> |
| 1997 | schizophrenia | Antipsychotic agents<br>Biperiden<br>Benserazide, Levodopa drug combination<br>Piribedil<br>Fluspirilene<br>Periciazine<br>Zopiclone<br>Promethazine<br>Venlafaxine hydrochloride<br>Prochlorperazine | Antiepileptic agents<br>Vigabatrin<br>Ethosuccinimide<sup>#</sup><br>Pokeweed*<sup>#</sup><br>25 hydroxycholecalciferol<br>5 HT uptake inhibitors<br>Tricyclic Antidepressive Agents<br>Anticonvulsive agents<br>Human insulin-like growth factor I<br>(1-34)-Human Parathyroid Hormone |

\*Diagnostic agents
<sup>#</sup>Drugs uncovered only through A→B→C association (see Table 2)

### Predictions prior to FDA approval

The FDA approval process can take several years. The manufacturers submit a new drug application or biologic license application, depending on the type of the product for which they seek approval. The average estimated time required by the FDA from phase zero of clinical trials to marketing approval is about 96.8 months (more than eight years)^37^. We evaluated SKiM on drugs approved by the FDA for migraine, AD, and schizophrenia.

The FDA has approved no drugs for RD, 22 drugs for migraine, six drugs for AD, and 11 drugs for schizophrenia^34^. To evaluate SKiM predictions, we collected the active ingredients of the approved drugs from the FDA and CenterWatch (https://www.centerwatch.com/), a source of information related to clinical trials for clinical research professionals. SKiM uncovered 12 drugs for migraine, five drugs for AD, and five drugs for schizophrenia at the cut-off date one year prior to the discoveries by Swanson and his colleagues. SKiM achieved 0.5641 recall on rediscovering active ingredients from FDA-approved drugs for migraine, AD, and schizophrenia (Supplementary Data 6). Manual analysis showed that 10 drugs for migraine and five drugs for schizophrenia are mentioned in PubMed only after the cut-off date, and thus these drugs could never be discovered by text mining on PubMed prior to the cut-off date. We excluded these drugs and calculated recall. SKiM achieved 0.9167 recall on active ingredients from FDA-approved drugs (Supplementary Data 6, Supplementary Note 9).

Manual analysis revealed SKiM predictions up to 32 years prior to approval by the FDA. In addition, we analyzed the predictions based on the validation by clinical trials, and the first publication on the therapeutic use of the drugs. SKiM predicted FDA-approved drugs up to 24 years prior to the validation by clinical trials, and up to 24 years prior to the first publication on the therapeutic use of the drugs (Supplementary Data 6). We believe that SKiM will be useful to clinicians, physicians, researchers, and pharmaceutical companies to accelerate drug repurposing.

### Finding new drug candidates for repurposing

We ran SKiM on PubMed abstracts published until March 2019 to look for new drug candidates for RD, migraine, AD, and schizophrenia. We evaluated the findings using DDA (positive controls) and manually analyzed the top 20 predictions to report the new potential drug candidates (Table 6). SKiM uncovered many new drugs from PubMed that are not in the DDA. The proportion of new candidate drugs for RD, migraine, AD, and schizophrenia indicates the wealth of information available in PubMed (Supplementary Table 3). The new drugs uncovered by SKiM may include drugs that may cause the disease of interest (negative associations). Though the current version of SKiM does not distinguish between the drugs useful for a disease and the drugs that may lead to the disease, our ranking approach typically places the most promising drugs at the top. While most of the negatively associated drugs rank lower in the list, a few are found among top 20 predictions: two drugs for RD, three drugs for migraine, and one drug for AD (Supplementary Data 7). In our experience, when we evaluate the top 20 predictions, negatively associated drugs are typically easily spotted.

**Table 6.**
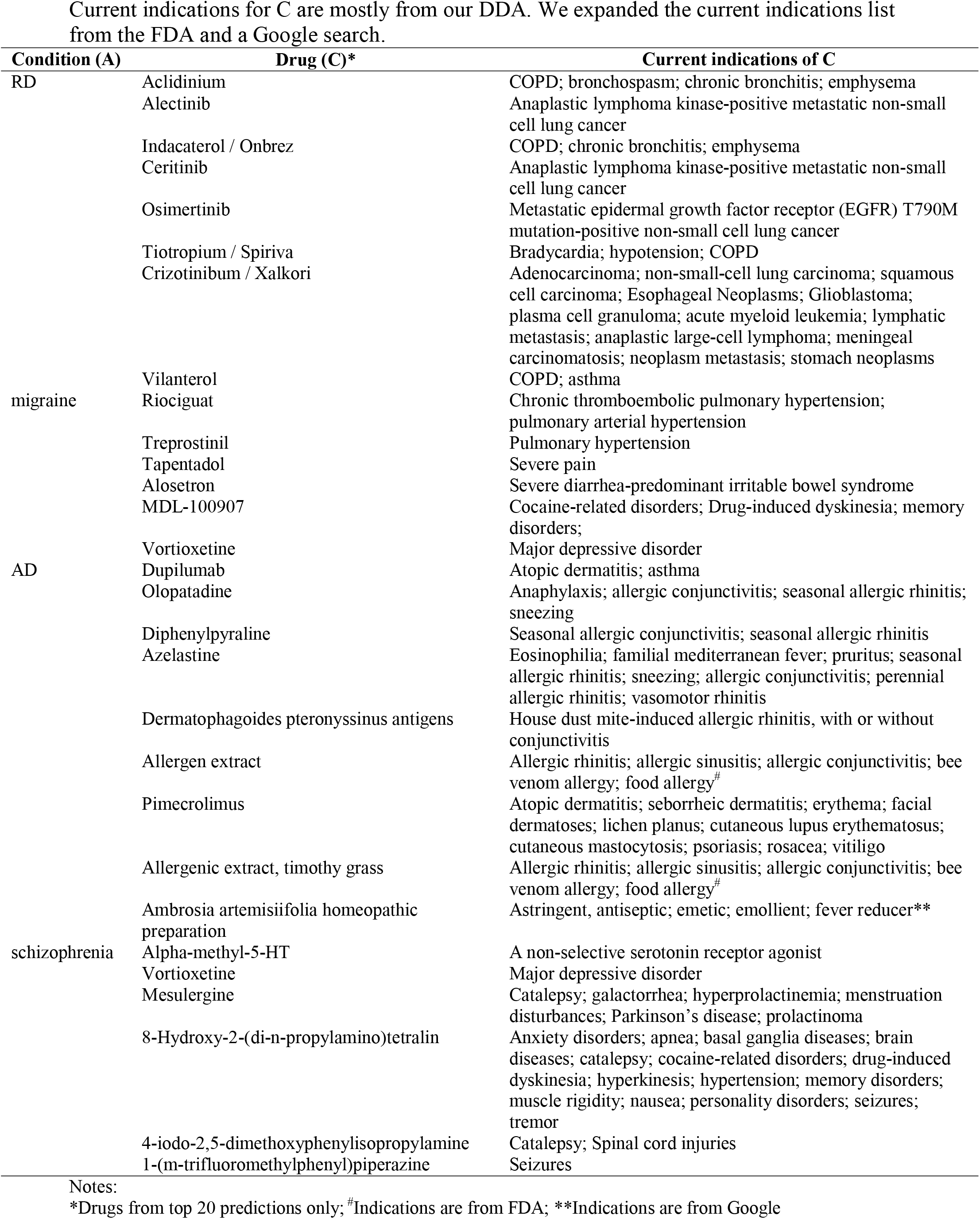
New potential drugs in top 20 predictions at cut-off date March 2019

The standard metrics such as precision and recall are not applicable for evaluating the new drugs, since “truth” about these new drugs with regard to new uses is not known. Alternatively, the prediction score (see Methods for details) of the annotated drugs from DDA and new drugs can be compared. Most of the new drugs are reported with a low prediction score when compared to the annotated drugs from DDA (Fig. 4). This indicates weak literature evidence on linking a new drug to a disease through phenotypes and symptoms. Certain new drugs are reported with a high prediction score (see “D” in Fig. 4). Manual annotation on the top 20 new drugs for each disease revealed many promising drug candidates for wet lab analysis and clinical trials (Table 6, Supplementary Data 7). Certain new drugs have evidence based on our literature analysis (Supplementary Data 7), but these drugs are missing in the DDA. Overall, SKiM achieved recall greater than 0.90 for RD, migraine, AD, and schizophrenia, and precision@20 is high enough (0.5882-0.9474) (see Supplementary Table 4 for details) that wet lab experiments on the top predictions listed in Table 6 would likely yield many positives. Thus, SKiM provides a short list of promising candidates for wet lab analysis. The drugs listed in Tables 2 and 6 are all candidates for further wet lab experiments and clinical trials.

**Fig 4.**
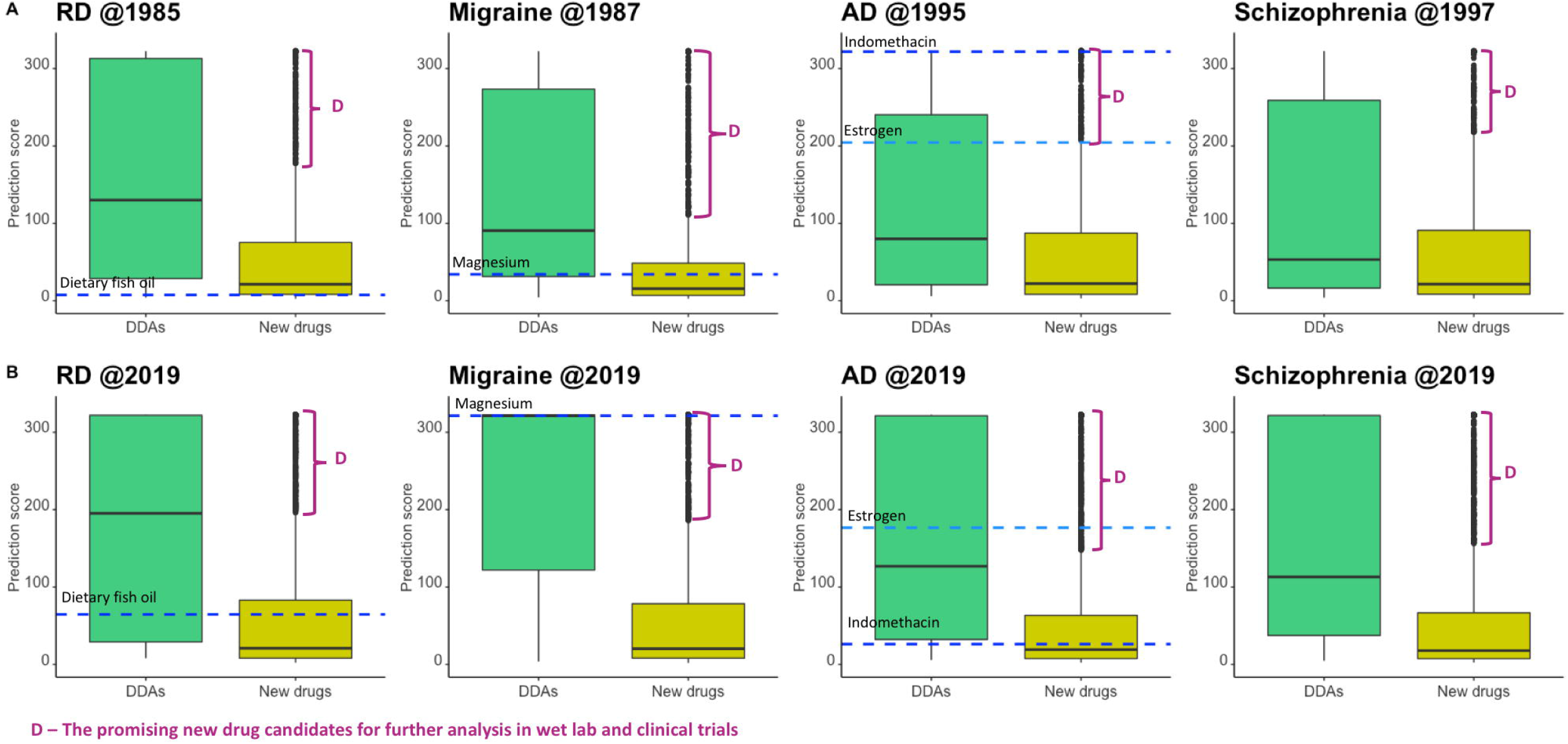
Drugs repurposed for RD, migraine, AD, and schizophrenia. **A.** Drugs repurposed for RD, migraine, AD, and schizophrenia at the cut-off date one year prior to the discoveries by Swanson and his colleagues. The drugs discovered by Swanson and his colleagues, dietary fish oil for RD, magnesium for migraine, and indomethacin and estrogen for AD are shown as dashed lines. The new drugs with high prediction score (“D”) are interesting ones for further study. The top 20 drugs from those with high prediction scores for each disease are listed in Table 2. Note that even when setting the cut-off date to 1985, 1987, or 1995, many new promising candidate drugs are suggested by SKiM. **B.** Drugs repurposed for RD, migraine, AD, and schizophrenia at the cut-off date March 2019. The drugs discovered by Swanson and his colleagues, dietary fish oil for RD, magnesium for migraine, and indomethacin and estrogen for AD are shown as dashed lines. The new drugs with high prediction score are interesting ones for further study. The top 20 drugs from those with high prediction scores for each disease are listed in Table 6. In **A** and **B**, the C term, calcium-independent phospholipase A2 (CIP-A2) from Swanson’s work for schizophrenia is not a drug but an enzyme associated with schizophrenia. Moreover, this is not a discovery in that the association between CIP-A2 and schizophrenia was well known before Swanson’s work^64^. Therefore, CIP-A2 is not highlighted.

### Repurposing drugs for 22 conditions

SKiM is a general LBD system applicable to repurposing drugs for any disease. In addition to four diseases from Swanson’s work, we also applied SKiM for repurposing drugs for 22 conditions with high prevalence rate (>200,000 cases each year) in the US and world-wide at cut-off date March 2019: type 2 diabetes (T2D), type 1 diabetes (T1D), atrial fibrillation, bipolar disorder, hypercholesterolemia, coronary heart disease (CHD), Grave’s disease, myocardial infarction, Parkinson’s disease, psoriasis, congestive heart failure (CHF), chronic obstructive pulmonary disease (COPD), emphysema, asthma, stroke, influenza, pneumonia, chronic kidney disease (CKD), kidney failure, suicide, lower respiratory tract infection (LRI), and tuberculosis.

We evaluated SKiM predictions using expert curated DDA for the 22 conditions (Supplementary Table 5) and compared the performance with KinderMiner. The performance of SKiM is better than KinderMiner (paired Wilcoxon test p-value = 4.768×10^-07^) (Supplementary Figure 2). While KinderMiner retrieved no DDA for CHD, suicide, and LRI, SKiM achieved 0.9130 recall for CHD, 1.0 recall for suicide, and 0.9677 recall for LRI. Precision@20 achieved by SKiM is low when evaluated using DDA (E1 in Supplementary Figure 3). Precision@20 significantly improved with manual analysis on finding literature evidence for A-C association and A-B-C association (E2 and E3 in Supplementary Figure 3). Thus, SKiM is applicable to any disease given that there is sufficient mention of the disease in the literature. SKiM uncovered many new drugs for 22 conditions. For any condition, the new drugs with the highest prediction scores are the interesting ones for further study (see D in Supplementary Figure 4, Supplementary Data 8). Our manual analysis on top 20 new drugs uncovered up to 15 new potential drugs for a disease (Supplementary Data 8). These are promising candidates for wet lab experiments and clinical trials. We discuss two conditions, T2D and suicide, in detail in the following two sections.

### Case study 1: Type 2 diabetes

Among 22 conditions, we chose T2D as a disease model to explore SKiM predictions. Diabetes is the seventh leading cause of death and the most expensive chronic condition in the US (total estimated cost of $327 billion every year)^14,30^. SKiM uncovered DDAs and new drugs for T2D (see T2D in Supplementary Figure 4). We reported the performance of SKiM on expert curated DDAs, and 25 drugs approved by American Diabetes Association (ADA)^38^. SKiM achieved 0.8242 recall when evaluated using expert curated DDAs and 1.0 recall on ADA-approved drugs. Precision@20 is 0.75 (see T2D in Supplementary Data 8). Among 25 ADA-approved drugs, 23 are DDAs. SKiM uncovered several new drugs with prediction scores higher than the remaining two drugs, ertugliflozin and bromocriptine (see T2D in Supplementary Figure 4). Manual analysis on the top 20 drugs confirmed four new potential drugs for T2D: riociguat, treprostinil, ambrisentan, and alirocumab (Supplementary Data 8).

### Case study 2: Suicide

The Center for Disease Control and Prevention reports a 33% increase in death rate due to suicide between the years 1999 and 2017 (https://www.cdc.gov/). Many biological, clinical, psychological, social, and environmental factors are leading causes for suicide^39^. Recent research on decreasing the death rate due to suicide focus on identifying the risk factors and their prevention^40,41^. Interestingly, ClinicalKey suggests two expert curated DDAs, brodalumab and tramadol for suicide. SKiM uncovered both the drugs and achieved 1.0 recall. SKiM also uncovered many new drugs for suicide (see suicide in Supplementary Figure 4). Manual analysis on the top 20 predictions showed six new potential drugs with literature evidence (see suicide in Supplementary Data 8). These drugs may be useful in decreasing the death rate due to suicide.

## Discussion

We describe SKiM, a generalized domain-independent open discovery LBD system. Unlike existing LBD systems, SKiM is capable of processing any combination of A, B, and C terms. SKiM links the knowledge across the abstracts to uncover both known and new discoveries. We tested SKiM on five discoveries by Swanson and his colleagues. SKiM uncovered all the discoveries at a default FET p-value less than 1×10^-05^. In the current study, we applied SKiM on repurposing drugs for four diseases from Swanson’s work: RD, migraine, AD, and schizophrenia, and 22 additional high-impact conditions. Using “time-slicing,” where we only allow SKiM to “read” abstracts from the first papers in PubMed (in the late 19th century) up to a particular cut-off date, we show that SKiM can uncover drugs useful for a disease up to 32 years before the first co-occurrence of a drug and a disease in a PubMed abstract, up to 16 years before the first mention of the usefulness of the drug in the literature, up to 31 years before the first validation by clinical trials, and up to 32 years before the approval by the FDA. Thus, SKiM uncovers potentially useful discoveries many years before they are directly mentioned in the literature or validated by clinical trials.

When SKiM was allowed to “read” all PubMed abstracts from the 19th century through March 2019, it uncovered almost twice the number of drugs for RD, migraine, AD, and schizophrenia than the time-slicing approach. Our manual analysis showed prediction of many new candidate drugs that have no direct literature evidence, but have evidence only via the LBD link of A-B and B-C. These are excellent new candidates for wet lab validation. Thus, SKiM is useful to narrow down the number of promising candidate drugs for follow up study.

Important for allowing SKiM to comprehensively scour the literature for associations, we compiled two comprehensive lexicons, a phenotypes and symptoms lexicon as B, and a drugs and biologics lexicon as C from various existing resources. Our findings are comprehensive and promising. For applying SKiM on a different application, similar lexicons are useful to uncover possible discoveries from PubMed abstracts. We compiled DDA from four expert curated resources, Comparative Toxicogenomics Database (CTD)^3^, National Drug File – Reference Terminology (NDF-RT)^42^, DrugBank^14^, and ClinicalKey (https://www.clinicalkey.com/info/healthcarefacilities/) for evaluating SKiM findings. The DDA includes therapeutically useful drugs for treating diseases. However, it lacks many therapeutically useful drugs published in PubMed abstracts. Many known drugs, such as cilostazol and vitamin D for RD, and vigabatrin for schizophrenia, are missing in the DDA. SKiM predicted these drugs at a cut-off date one year prior to the discoveries by Swanson and his colleagues and advanced the discovery up to 28 years prior to their first publication that indicates the usefulness of a drug for a disease: cilostazol by 18 years^43^, vitamin D by 28 years^44^, and vigabatrin by five years^45^. Recently, cilostazol^46^ and vitamin D^44^ for RD were validated by clinical trials. Thus, SKiM is useful for extracting information from PubMed abstracts that is yet to be explored by scientists.

SKiM uncovers both expert curated drugs from DDA and new drugs for RD, migraine, AD, and schizophrenia. Interestingly, SKiM repurposed certain drugs from DDA to a new disease: adofen, a known drug for RD is repurposed for migraine; and atenolol, a known drug for migraine is repurposed for RD. SKiM repurposed adofen for migraine five years prior to validation by clinical trials, and atenolol for RD four years prior to validation by clinical trials. The repurposed drugs, adofen for migraine and atenolol for RD, rank among the top 20 predictions (Supplementary Data 3). Thus, SKiM is reliable for repurposing drugs for various diseases.

For existing LBD systems, co-occurrence approaches tend to produce a large number of false positives (FP). In SKiM, we partially address the issue by filtering statistically significant A➔B and B➔C pairs and present a sorted list of C terms. Similar to existing LBD systems, SKiM uses top n=50 B terms from A➔B and processes C terms for every B term. It is possible to miss certain C terms associated only with B terms that are not among the top 50 predictions (see Supplementary Discussion 1).

Using SKiM, we predicted new potential repurposable drugs for RD, migraine, AD, and schizophrenia, and 22 additional conditions with a high prevalence rate. SKiM can be applied to repurpose drugs for any disease that has sufficient information in PubMed. Furthermore, SKiM is a generalized LBD system, and it should be applicable to a variety of inquiries about biomedicine, not just drug repurposing. The approach is simple, yet powerful. We believe that SKiM as a generalized tool will support a wide range of researchers by allowing them to focus their experimentation on a short list of targets. To our knowledge, SKiM is the only reliable, functioning generalized system for drug repurposing using an LBD approach. The code and instructions needed for building an indexed PubMed corpus and running SKiM are available here (https://github.com/stewart-lab/SKiM).

### Future work

No text mining system is perfect^2^. SKiM findings include FPs. The pharmacological effect of a drug co-occurring with a disease in a PubMed abstract may indicate either treatment or cause of the disease. The current version of SKiM does not differentiate between positive and negative associations. Our manual analysis revealed a few FPs among top 20 predictions for RD, migraine, AD, and schizophrenia (Supplementary Data 3). Fortunately, FPs are typically easily spotted by manual analysis. In the future, we will address the limitations of SKiM by using advanced text mining approaches such as BioBERT^47^, ELMo^48^, CoVe^49^, or other methods to reduce FPs. These advanced approaches are capable of distinguishing positive associations (i.e. the drug is useful in treating a disease) from negative associations (i.e. the drug may cause the disease).

## Methods

### Overview

The approach consists of the following steps: (1) building PubMed abstracts (P) with abbreviations replaced to original text; and (2) querying P with an initial concept A to retrieve a set of C concepts, through a set of B concepts (i.e. A➔B➔C). We applied SKiM on repurposing drugs for RD, migraine, AD, and schizophrenia, and 22 additional conditions with a high prevalence rate. We used phenotypes and symptoms as B, and drugs as C. We evaluated the predictions by manual analysis and found many of the predictions can be made using only literature available from many years before their validation by clinical trials and approval by the FDA.

### PubMed repository

The Center for High Throughput Computing (CHTC) at the University of Wisconsin-Madison (http://chtc.cs.wisc.edu/) maintains ~31 million PubMed abstracts to support 296 research projects, including SKiM. CHTC updates the resource quarterly by downloading PubMed baseline XML files and applying daily updates from the FTP server maintained by the National Library of Medicine (https://www.nlm.nih.gov/). Almost every PubMed abstract contains abbreviations, which are highly ambiguous^50^. We replace abbreviations with the original text using Allie^51^, a database of PubMed ID (PMID), abbreviations, and the original text from PubMed abstracts. Allie is released every month. CHTC maintains two versions of PubMed abstracts, one with abbreviations to support the existing projects and the other with abbreviations replaced to the original text to exclusively support SKiM. Both versions are accessible to SKiM through distinct field names. In the current study, we used 29,613,663 PubMed abstracts (downloaded in March 2019) with 76,904,011 abbreviations replaced to the original terms.

### KinderMiner and Query improvements

Recently we developed a simple text mining system called KinderMiner for querying PubMed abstracts. The system filters and sorts a list of target terms (e.g. magnesium) by their association with a key phrase (e.g. migraine) based on co-occurrence significance and proportion^19^. Querying key phrase or target term in the article title or abstract is an efficient approach because not all articles contain an abstract (e.g. PMID: 30561177, 30586516). Both the key phrase and target term may include synonyms, and their identification is mandatory to get the accurate PubMed abstracts counts for statistical analysis. We updated the KinderMiner algorithm to perform a Boolean search of key phrase, target term, and the synonyms in the article title or abstract:

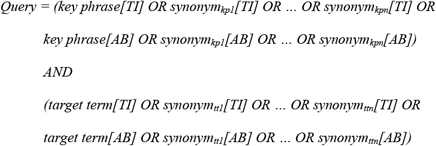

where ‘*TI*’ refers to the article title and ‘*AB*’ refers to the abstract. The suffixes ‘*kp*’ refers to key phrase and ‘*tt*’ refers to target term. The key phrase in the query is ignored to get the number of PMIDs with target term (*N*_*t*t_). The target term in the query is ignored to get the number of PMIDs with key phrase (*N_kp_*). Presence of both key phrase and target term in the query gets the number of PMIDs with key phrase and target term (*N*_*k*t_). The query can be pointed to PubMed repository with abstracts with abbreviations, or abstracts with abbreviations replaced to the original text. We released the updated algorithm as KinderMiner 2.0 (https://github.com/stewart-lab/KinderMiner_2).

### Mining known and new knowledge

SKiM is meant for querying PubMed in a serial manner (Fig. 1). SKiM uses the features of KinderMiner for retrieving A➔Bs and Bs➔Cs. Our recent work on retrieving mitochondrial PPI from PubMed using KinderMiner 2.0 performed better than PolySearch2^2^ and STRING, a popular database on PPI^23^. For a given input A, the query retrieved a list of significant Bs cooccurring with A. The significant Bs were filtered on an arbitrary FET p-value less than 1×10^-05^ and sorted on the prediction score, which is calculated as the logarithmic sum of FET p-value and sort ratio of B➔Cs to get the known discoveries at the top^19^.

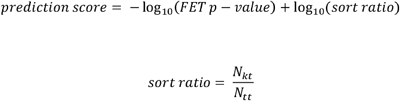

Here, *N_kt_* represents the number of PMIDs with co-occurring A and B, or B and C. *N_tt_* represents the number of PMIDs with B term in the case of A➔B, and represents the number of PubMed abstracts with C term in the case of B➔C. Similar to existing LBD systems^8,52,53^, we select the top N significant Bs for retrieving significant Cs, where N is the count of top results kept. For every significant B from top N, the query retrieved a list of significant Cs that cooccurred with B. All Bs➔Cs are combined and sorted on the prediction score. SKiM reports every B➔C with the corresponding prediction score. Note that any particular C may co-occur with multiple Bs. We performed a second sorting of Bs➔Cs. For each C, we retrieved all Bs and sorted on the prediction score. Our ranking approach is designed to push the known Cs to the top of the results. In the current study, we do not consider the ranking order of A➔Bs for Bs➔Cs ranking.

SKiM provides a variety of options to query PubMed. We adopted three options from KinderMiner: *(i)* querying PubMed with tokenized A, B, and C, *(ii)* querying PubMed with stemmed A, B, and C, and *(iii)* querying PubMed abstracts published until a preferred year (i.e. cut-off date). KinderMiner provides tokenization and stemming for the key phrase alone. SKiM extends the tokenization and stemming for A, B, C terms and their synonyms. The default is to query PubMed without tokenization and stemming.

### Drug repurposing application

SKiM is a general LBD system for discovering something about A, where A can be a biomedical concept such as disease, gene, or a pathway. B and C are a list of concepts from which SKiM uncovers the most significant Cs for A through Bs. In the current study, we applied SKiM on repurposing drugs for RD, migraine, AD, and schizophrenia, and 22 additional high-impact conditions. In the A➔Bs➔Cs work-flow, A refers to a disease, B refers to the list of phenotypes and synonyms, and C refers to the list of drugs. For us to perform these searches, we first constructed the lexicon for each category of terms.

#### Diseases lexicon

We compiled a diseases lexicon from UMLS Metathesaurus^13^ and SNOMED CT^54^. The 2018AB version of UMLS Metathesaurus was installed in Rich Release Format (RRF) using MetamorphoSys, a UMLS installation wizard. The resource contains 3.64 million health related concepts (or 13.9 million unique concepts names) from 201 source vocabularies (e.g. MeSH, RxNorm). Within UMLS Metathesaurus, we used MRCONSO, a resource on the concepts and MRSTY, a resource on the sematic types for the concepts. Though UMLS Metathesaurus includes a complete version of SNOMED CT, the default installation includes only 54 SNOMED concepts. SNOMED CT within UMLS Metathesaurus is installed by selecting a specific edition (i.e. SNOMED CT US Edition) during installation using MetamorphoSys. This version of SNOMED CT includes unique concept identifier (CUI) from UMLS Metathesaurus, and we assigned the semantic types to SNOMED concepts using CUIs.

We filtered the concepts belonging to semantic types under “Disorder”: UMLS Metathesaurus consists of 163,073 diseases (or 437,580 diseases and synonyms), and SNOMED CT consists of 146,436 diseases (or 470,076 diseases and synonyms). We combined the resources and removed the duplicates and 84 disease synonyms matching with English stop-words (e.g. in, present). The process obtained 273,710 diseases (or 640,386 diseases and synonyms). 7,450 diseases and synonyms include more than one CUI in UMLS Metathesaurus. For example, T2D and its synonyms are assigned with two CUIs, C0011860 and C4014362. We combined diseases and synonyms with more than one CUI and assigned a customized CUI which starts with ‘M’ to represent modified ‘CUI’. All the semantic types from multiple CUIs are assigned to the modified CUI. Our disease lexicon consists of 264,983 diseases (or 640,386 diseases and synonyms).

#### Phenotypes and symptoms lexicon

We compiled a phenotypes and symptoms lexicon from Human Phenotype Ontology (HPO)^55^, Phenome Wide Association Studies (PheWAS)^56^ and Online Mendelian Inheritance in Man (OMIM)^57^ (Supplementary Figure 5). We downloaded the hp.obo file from http://human-phenotype-ontology.github.io/ and extracted 29,263 phenotypes and synonyms. We downloaded 414 phenotypes from OMIM (https://www.omim.org/phenotypicSeriesTitle/all/) and 1,866 phenotypes from PheWAS catalog (https://phewascatalog.org/phecodes). The identifiers in HPO, OMIM, and PheWAS catalog are resource specific. We assigned a customized identifier and removed the duplicates across the resources. The process obtained 31,156 phenotypes and synonyms. None of the resources include blood viscosity from Swanson’s discovery. HPO contains ‘blood hyperviscosity’ and two synonyms, ‘hypercoagulability’ and ‘thrombophilia’. We extended the phenotypes and synonyms list by deriving new terms from the existing ones (i.e. ‘blood hyperviscosity’ to ‘blood viscosity’). We used a list of 500 prefixes and suffixes related to the medical terminology^58,59^ to remove prefix/suffix from phenotypes and synonyms. Our approach obtained 90,634 new terms. However, all the new terms are not phenotypes or symptoms (e.g. ‘plastic kidney’ from ‘dysplastic kidney’, ‘opathy’ from ‘nephropathy’, and ‘renal s’ from ‘renal cysts’). We used KinderMiner to filter 3,314 new terms mapping to at least two PubMed abstracts, and manually curated 1,771 new phenotypes that are not in HPO, OMIM, and PheWAS. We used the diseases lexicon to assign CUI to phenotypes and symptoms. All the phenotypes and symptoms do not map to CUI from the diseases lexicon. We assigned customized CUIs for those without a CUI. For every phenotype or symptom, we included all the synonyms from the diseases lexicon. The approach obtained a phenotypes and symptoms lexicon with 35,229 phenotypes, symptoms, and their synonyms (i.e. 9,272 phenotypes and symptoms).

#### Drugs lexicon

Our drugs lexicon^16^ is from UMLS Metathesaurus 2018AB version^13^, DrugBank 5.0^14^, and PharmGKB^15^. In brief, we filtered the drugs and biologics from UMLS Metathesaurus based on four semantic types: clinical drug, antibiotic, pharmacologic substance, and immunological factor, and included the missing drugs and synonyms from DrugBank and PharmGKB. We assigned customized drug ID instead of CUI to accommodate drugs from DrugBank and PharmGKB that are not in UMLS Metathesaurus. The customized drug ID includes a prefix ‘CD’ (e.g. CD00014279 for zotepine). Similar to diseases lexicon, we combined the drugs and synonyms with multiple customized drug IDs and assigned new customized drug IDs with ‘CDM’ prefix, where ‘M’ stands for modified (e.g. CDM00000663 for metformin). The process obtained 693,314 drugs and synonyms.

#### Drugs subset for repurposing application

The drugs lexicon has a plethora of general drug classes (e.g. antihypertensive agents), general concepts (e.g. antigens), and pharmacologic substances that are not drugs (e.g. chemical buffer). For drug repurposing, we derived a subset of drugs associated with disease, gene, phenotype, variant, or haplotype from various resources. We retrieved 5,288 drugs associated with diseases from CTD^3^, NDF-RT^42^, DrugBank^14^, and FDA^34^. Many FDA-approved drugs are missing in CTD, NDF-RT, and DrugBank, and vice-versa. We retrieved 7,293 drugs associated with genes from CTD, DrugBank, and PharmGKB^15^. We retrieved 2,620 drugs associated with phenotypes from CTD. We retrieved 642 drugs associated with variants or haplotypes from PharmGKB. We included 3,080 annotated drugs from PharmGKB. The drugs from various associations and annotations overlap (Supplementary Figure 6). We utilized the drugs lexicon to remove the overlapping duplicates across the resources by mapping the drugs to drug ID in the drugs lexicon. The process obtained 9,665 drugs.

### Evaluating SKiM on existing approaches

One common method for evaluation of LBD systems^10,60,61^ is to assess how well the LBD system rediscovers the five discoveries by Swanson and his colleagues. The approach is simple: the system is allowed to query the A term from the discovery (e.g. RD) on PubMed abstracts published through a cut-off date before the discovery (e.g. the year, 1985). The system is expected to predict the associated C term (e.g. dietary fish oil). LBD systems are also evaluated on predicting a discovery prior to the first landmark publication validating the discovery in PubMed. We evaluated SKiM on rediscovering the discoveries by Swanson and his colleagues from PubMed articles published until one year before the discovery. We evaluated top 20 SKiM findings manually using PubMed, clinical trials, and the FDA to find the number of years SKiM can advance a discovery. In addition, we also evaluated SKiM based on a recent work on automated time-slicing (Supplementary Method 2 and Supplementary Method 2 Table 3)^62^.

### Datasets used for evaluating SKiM for drug repurposing

The DDA is compiled from four resources, Comparative Toxicogenomics Database (CTD)^3^, National Drug File – Reference Terminology (NDF-RT)^42^, DrugBank^14^, and ClinicalKey (https://www.clinicalkey.com/info/healthcarefacilities/). CTD includes both expert curated drugs and inferred drugs for various diseases. We filtered the expert curated drugs annotated for ‘therapeutic use’ from CTD. We used the disease-drug association list compiled recently from CTD and NDF-RT^63^ and extended it with the disease-drug association from DrugBank and ClinicalKey. DrugBank includes the disease-drug association in the indications section. We downloaded DrugBank in XML format and retrieved 3,108 indications. We manually curated the associations for RD, migraine, AD, and schizophrenia, and 22 additional high-impact conditions. We excluded the drugs that are currently under investigation for a disease. We manually queried the web interface of ClinicalKey to retrieve the drugs suggested for RD, migraine, AD, and schizophrenia, and 22 additional high-impact conditions. The drug identifiers across resources are different. We assigned the identifier from our drugs lexicon and removed the duplicates. Finally, we eliminated the drug combinations from our collection because our objective is to repurpose individual drugs (Supplementary Table 2, Supplementary Table 5).

## Acknowledgments

KR, JS, FK, JT, and RS acknowledge funding from the Morgridge Institute for Research and a grant from Marv Conney. IR acknowledges the GeoDeepDive Infrastructure, funded by NSF ICER 1343760. LCT is supported by the National Psoriasis Foundation, award from the National Institutes of Health (K01AR07212), as well as the Babcock Memorial Trust. ZN acknowledges funding from the National Institutes of Health (NIH GM102756).

## Contributions

K.R. and R.S. designed the study and evaluation. K.R. and J.S. developed KinderMiner 2.0 and SKiM. I.R. and M.L. built the local PubMed repository. I.R. and K.R. contributed to integrate Allie with PubMed repository for handling abbreviations. K.R., L.C.T., and R.S. compiled the lexicons from various existing resources. K.R., R.S., Z.N., F.K., and L.C.T. provided statistical support. J.T. provided biological inferences or interpretation of the results. K.R. and R.S. wrote the manuscript, and every author has reviewed the work.

## Competing interests

None.

## Code availability

The code for SKiM is available on GitHub (https://github.com/stewart-lab/SKiM). SKiM needs a local version of PubMed repository. We share the code for generating the repository in GitHub (https://github.com/iross/km_indexer). The code for KinderMiner 2.0 is available in GitHub (https://github.com/stewart-lab/KinderMiner_2). In addition, we also provide the code for compiling various lexicons for drugs repurposing application in GitHub. The code for compiling diseases lexicon is at https://github.com/stewart-lab/Diseases_lexicon. The code for compiling phenotypes and symptoms lexicon is at https://github.com/stewart-lab/Phenotypes_and_symptoms_lexicon. The code for compiling drugs lexicon is at https://github.com/CutaneousBioinf/LiteratureMiningTool/tree/master/DrugDict. Code for retrieving the drugs subset from drugs lexicon is at https://github.com/stewart-lab/Drugs_subset_for_drugs_repurposing_application. The code for compiling expert curated disease-drug associations (DDA) from CTD and NDF-RT is at https://github.com/CutaneousBioinf/LiteratureMiningTool/tree/master/RelatGold/Disease2DrugAssociationGoldStandard. The code for processing DDA from DrugBank and ClinicalKey, and to combine DDAs from CTD, NDF-RT, DrugBank, and ClinicalKey is at https://github.com/stewart-lab/Expert_curated_disease_drug_associations. The code for simulation study to find false discovery rate (FDR) is at https://github.com/zijianni/SKiM_simulation.

## Notes

### Competing Interest Statement

The authors have declared no competing interest.

