## Supplementary File for "SKiM - A generalized literature-based discovery system for uncovering novel biomedical knowledge from PubMed"

Home page: https://morgridge.org/profile/ron-stewart/

**Supplementary Notes**

**Supplementary Note 1:** SKiM predicted “blood viscosity” as a B term for uncovering dietary fish oil for Raynaud’s disease (RD). Interestingly, “blood viscosity” is one of the B terms reported by Swanson for the same discovery^1^. Swanson and his colleagues identified drug, hormone, inflammation, and other concepts as B terms for the remaining four discoveries^2^. SKiM rediscovered all five discoveries by Swanson and his colleagues by using phenotypes and symptoms as B terms (Supplementary Data 1). Thus, the phenotypes and symptoms as B terms are appropriate for drug repurposing and will likely be useful for other tasks. We are not including Swanson’s work on the association of schizophrenia and calcium-independent phospholipase A2 (CIP-A2)^3^ as a discovery because there are 25 papers on schizophrenia and CIP-A2 association before his publication.

**Supplementary Note 2:** BITOLA has an online functional interface (http://ibmi.mf.uni-lj.si/sl/bitola). The tool is restricted to select only UMLS concepts as A, B, and C terms. BITOLA does not provide an option to define a cut-off date. The default cut-off date is unknown. For these reasons, BITOLA cannot be compared with SKiM using the same settings.

We explored BITOLA with the default settings on rediscovering the five discoveries by Swanson and his colleagues over the years. BITOLA uncovered three discoveries using Disorders or UMLS concepts as B terms (see Supplementary Note Table 1). For discoveries on migraine and Alzheimer’s disease (AD), BITOLA uncovered C terms containing magnesium and estrogen, not the exact C term defined in the discoveries. We also noticed that BITOLA does not show the intermediate B terms connecting A and C terms. Though BITOLA has a functioning user interface, it does not allow the use to provide a cut-off date to validate the findings on time-slicing, an existing evaluation approach suggested for LBD systems.

Supplementary Note Table 1. Performance of BITOLA on Swanson’s discoveries

| **A** | **C** | **Rediscovery using BITOLA** | |
| --- | --- | --- | --- |
|  |  | **Disorders as B** | **UMLS concepts as B** |
| RD | Dietary fish oil | Yes | Yes |
| migraine | Magnesium | No | No |
| AD | Indomethacin | Yes | Yes |
| AD | Estrogen | No | No |
| Somatomedin C | Arginine | Yes | Yes |

BITOLA is set to display top 50 B terms from A→B and top 100 C terms from B→C. Similar to SKiM, we selected top 50 B from A→B to uncover C terms. The number of B terms uncovered by BITOLA from A→B is listed in Supplementary Note Table 2. The number of B terms uncovered by SKiM at cut-off date one year prior to the discoveries by Swanson and his colleagues and at cut-off date March 2019 to include all PubMed articles are listed in Supplementary Table S3. Though we cannot compare the number of B terms uncovered by BITOLA with the number of B terms uncovered by SKiM, it is obvious that the number of B terms uncovered by SKiM at cut-off date March 2019 is much less than the number of B terms uncovered by BITOLA at an unknown default cut-off date (Supplementary Note Table 2, Supplementary Note Table 3).

Supplementary Note Table 2. B terms connecting A and C terms in BITOLA

| **A** | **C** | **Cut-off date** | **B terms count** | |
| --- | --- | --- | --- | --- |
|  |  |  | **Disorders as B** | **UMLS concepts as B** |
| RD | Dietary fish oil | Unknown | 1,090 | 4,954 |
| migraine | Magnesium | Unknown | 1,457 | 7,772 |
| AD | Indomethacin | Unknown | 1,269 | 16,543 |
| AD | Estrogen | Unknown | 1,269 | 16,543 |
| Somatomedin C | Arginine | Unknown | 152 | 2,485 |

Supplementary Note Table 3. B terms connecting A and C terms in SKiM

| **A** | **C** | **B terms connecting A and C** | | | |
| --- | --- | --- | --- | --- | --- |
|  |  | **Cut-off date (Y1)** | **B terms count at Y1** | **Cut-off date (Y2)** | **B terms count at Y2** |
| RD | Dietary fish oil | 1985 | 101 | March 2019 | 287 |
| migraine | Magnesium | 1987 | 115 | March 2019 | 465 |
| AD | Indomethacin | 1995 | 233 | March 2019 | 399 |
| AD | Estrogen | 1995 | 233 | March 2019 | 399 |
| Somatomedin C | Arginine | 1989 | 81 | March 2019 | 143 |

**Supplementary Note 3:** Like BITOLA, LION LBD has an online functional interface (https://lbd.lionproject.net/). LION LBD is restricted to use only chemicals, diseases, mutations, genes, cancer hallmarks and species as A, B, and C terms. Dietary fish oil is not within the scope of the concept types used in LION LBD and thus Swanson’s discovery on RD cannot be tested^4^.

LION LBD is evaluated on four of five discoveries by Swanson and his colleagues at cut-off date five years prior to the discovery^4^ and its performance is unclear. In order to compare LION LBD with SKiM, we tested the open discovery system of LION LBD on five discoveries by Swanson and his colleagues at cut-off date one year prior to the discovery. LION LBD does not show C terms from the discoveries by Swanson and his colleague with default weight settings. The online interface of LION LBD does not provide an option to view more or all C terms uncovered for a given A term. With the online LION LBD interface, we are unable to recover any of Swanson and his colleagues’ discoveries. We also observed that LION LBD does not handle specific entity type (e.g. chemical, disease) as B or C terms.

**Supplementary Note 4:** We compiled expert curated disease-drug associations (DDA) from four existing resources, Comparative Toxicogenomics Database (CTD)^5^, National Drug File – Reference Terminology (NDF-RT)^6^, DrugBank^7^, and ClinicalKey (https://www.clinicalkey.com/info/healthcarefacilities/). We used DDA to evaluate the drugs repurposed by SKiM for Raynaud’s disease (RD), migraine, Alzheimer’s disease (AD), and schizophrenia.

SKiM achieved 0.8333 recall for RD, 0.7528 recall for migraine, 0.7049 recall for AD and 0.7935 recall for schizophrenia at the cut-off date one year prior to the discoveries by Swanson and his colleagues. SKiM predicted all or most of the drugs from DDA when the cut-off date is relaxed to March 2019, (achieving recall of 1.0 for RD, 0.9775 for migraine, 0.9180 for AD, and 0.9239 for schizophrenia). The recall achieved by SKiM is higher than the recall achieved by KinderMiner at both cut-off dates (Fig. 3A). Thus, SKiM (A🡪Bs🡪Cs) is capable of uncovering undiscovered public knowledge from PubMed more effectively than KinderMiner (A🡪Cs).

**Supplementary Note 5:** The precision achieved by SKiM and KinderMiner using the expert curated disease-drug associations (DDA) is very low (Table 1). This is expected because both the systems retrieve all possible drugs for a given disease. All the expert curated resources involve manual annotation of drugs useful for diseases. The process is tedious and time consuming, and cannot be effectively maintained with the large number of articles continually being added to PubMed. The knowledge gap between DDA and information in PubMed abstracts is huge.

In text mining, a common approach to evaluate systems such as SKiM is to calculate precision of top N predictions, where N is the count of included predictions (i.e. precision@N)^8^. For SKiM, we rank Cs such that the most relevant ones are at the top. To our surprise, precision@20 was low: 0.05 for RD, 0.05 for migraine, 0.00 for AD and 0.10 for schizophrenia using the DDA. This largely represents a deficiency in the existing DDA list. Our manual analysis revealed that many drugs known for treating the diseases in the literature are missing in the DDA (Supplementary Data 3). The precision@20 significantly improved after manual curation: 0.5882 for RD, 0.90 for migraine, 0.7895 for AD and 0.85 for schizophrenia (Fig. 3B, Table 1).

**Supplementary Note 6:** Some of the expert curated disease-drug associations (DDA) for Raynaud’s disease (RD), migraine, Alzheimer’s disease (AD) and schizophrenia are suggested for two or three diseases under study. For example, the drug Fluoxetine is curated for RD and schizophrenia (Supplementary Data 4). For evaluating SKiM on drugs annotated for more than one disease, we obtained a list of 60 pairs of disease and drug (Supplementary Data 4). SKiM uncovered 56 disease-drug pairs at cut-off date one year prior to the discoveries by Swanson and his colleagues: 1985 for RD, 1987 for migraine, 1995 for AD and 1997 for schizophrenia.

**Supplementary Note 7:** The discoveries by Swanson and his colleagues, dietary fish oil for RD^1^, magnesium for migraine^2^, and indomethacin and estrogen for AD^9,10^ were later successfully validated through clinical trials^11-14^. SKiM uncovered these discoveries by reading PubMed articles published until one prior to the discovery by Swanson and colleagues (Fig. 2B).

**Supplementary Note 8:** Metformin is one among the eight drugs that is not yet recognized as an expert curated disease-drug association for AD (Table 4). Interestingly, SKiM repurposed metformin, an antidiabetic drug for AD from PubMed articles published until 1995 (Table 4). 22 years later, the drug was validated for AD by clinical trials^15^. Recently, metformin has been validated by clinical trials for longevity in a mouse model^16^ and prostate cancer in men^17^. Thus, one drug might be useful for treating multiple conditions, and SKiM is an effective way to explore drugs useful for treating any condition of interest.

**Supplementary Note 9:** SKiM uncovered most of the FDA approved drugs for migraine AD and schizophrenia, and achieved 0.5641 recall. Certain drugs such as Excedrin Migraine and Namzaric approved by the FDA include more than one active ingredient. SKiM uncovered all the active ingredients for such drugs (Supplementary Data 6). The results show the wealth of information available in PubMed. LBD systems like SKiM can uncover many promising drugs for various diseases.

SKiM uncovered all FDA approved drugs for AD and schizophrenia when the cut-off date is relaxed to March 2019. It failed to predict four FDA approved drugs for migraine: Aimovig, Ajovy, Emgality, and Amerge. SKiM achieved recall of 0.8974 at a cut-off date of March 2019 (Supplementary Data 6).

We observed that 84.62% (33 out of 39) of drugs approved by the FDA are expert curated disease-drug associations (DDAs). Six drugs approved by the FDA are not among DDAs: naratriptan hydrochloride, almotriptan malate and eletriptan hydrobromide for migraine, donepezil hydrochloride for AD, and aripiprazole lauroxil and ziprasidone mesylate for schizophrenia (Supplementary Data 6). Thus, DDA does not include all the drugs approved by the FDA.

**Supplementary Tables**

**Supplementary Table 1:** Performance of KinderMiner on six discoveries by Swanson and his colleagues using PubMed abstracts published through March 2019

| **A** | **C** | **FET p-value**  **(A-C)** |
| --- | --- | --- |
| RD | dietary fish oil | 0.0187 |
| migraine | magnesium | 7.52×10^-53^* |
| AD | indomethacin | 0.9992 |
| AD | estrogen | 1.22×10^-41^* |
| somatomedin C | arginine | 8.02×10^-175^* |

*KinderMiner predicted the discovery at FET p-value less than1×10^-05^

**Supplementary Table 2.** Expert curated disease-drug associations (DDA) from various resources

| **Expert curated disease** | **Expert curated drugs / drug combinations** | | | **DDA** | **Expert curated drug combinations** |
| --- | --- | --- | --- | --- | --- |
|  | **CTD + NDF-RT** | **DrugBank** | **ClinicalKey** |  |  |
| RD | 13 | 6 | 1 | 18 | 1 |
| migraine | 58 | 24 | 60 | 89 | 18 |
| AD | 54 | 17 | 9 | 61 | 4 |
| schizophrenia | 80 | 38 | 22 | 92 | 2 |

**Supplementary Table 3.** Proportion of new drugs uncovered by SKiM

| **Disease** | **Cut-off date** | **Number of new drugs (N1)** | **Proportion (N1 / total drugs)** | **Cut-off date** | **Number of new drugs (N2)** | **Proportion (N2 / total drugs)** |
| --- | --- | --- | --- | --- | --- | --- |
| RD | 1985 | 1,604 | 0.1660 | March 2019 | 3,615 | 0.3740 |
| migraine | 1987 | 1,469 | 0.1520 | March 2019 | 3,325 | 0.3440 |
| AD | 1995 | 2,667 | 0.2759 | March 2019 | 3,452 | 0.3572 |
| schizophrenia | 1997 | 1,703 | 0.1762 | March 2019 | 2,509 | 0.2596 |

N1 and N2 are the number of new drugs uncovered by SKiM with cut-off date one year prior to the discoveries by Swanson and his colleagues and the year 2019 respectively. The proportion indicates the portion of the drugs repurposed for RD, migraine, AD and schizophrenia at the respective cut-off dates.

**Supplementary Table 4.** Recall and precision@20 of drugs uncovered by SKiM at cut-off date 2019

| Disease | All drugs | | | Top 20 drugs | | | |
| --- | --- | --- | --- | --- | --- | --- | --- |
|  | DDA only | | | DDA + Manual analysis | | | New potential drugs* |
|  | TP | FN | Recall | TP | FP | Precision@20 |  |
| RD | 18 | 0 | 1.0000 | 14 | 5 | 0.7368 | 8 |
| migraine | 87 | 2 | 0.9775 | 15 | 5 | 0.75 | 6 |
| AD | 56 | 5 | 0.9180 | 10 | 7 | 0.5882 | 9 |
| schizophrenia | 85 | 7 | 0.9239 | 18 | 1 | 0.9474 | 6 |

*Drugs uncovered only through A-B-C

**Supplementary Table 5.** Experts curated disease-drug associations (DDAs) for 22 conditions

| **Expert curated condition** | **Expert curated drugs / drug combinations** | | | | **DDA** | **Expert curated drug combinations** |
| --- | --- | --- | --- | --- | --- | --- |
|  | **CTD + NDF-RT** | **DrugBank** | **ClinicalKey** | **All** |  |  |
| type 2 diabetes (T2D) | 64 | 44 | 8 | 95 | 91 | 4 |
| type 1 diabetes (T1D) | 10 | 10 | 18 | 26 | 21 | 4 |
| atrial fibrillation | 53 | 22 | 29 | 70 | 70 | 0 |
| bipolar disorder | 78 | 12 | 19 | 86 | 83 | 3 |
| hypercholesterolemia | 60 | 17 | 34 | 79 | 68 | 11 |
| coronary heart disease (CHD) | 45 | 4 | 15 | 50 | 50 | 0 |
| Graves’ disease | 9 | 2 | 1 | 9 | 9 | 0 |
| myocardial infarction | 142 | 41 | 69 | 182 | 167 | 15 |
| Parkinson’s disease | 60 | 32 | 22 | 77 | 71 | 6 |
| psoriasis | 55 | 28 | 62 | 101 | 95 | 6 |
| congestive heart failure (CHF) | 50 | 35 | 58 | 114 | 100 | 14 |
| chronic obstructive pulmonary disease (COPD) | 31 | 13 | 48 | 74 | 61 | 13 |
| emphysema | 9 | 15 | 14 | 31 | 27 | 4 |
| asthma | 74 | 45 | 50 | 112 | 101 | 11 |
| stroke | 54 | 21 | 34 | 88 | 80 | 8 |
| influenza | 10 | 5 | 31 | 40 | 18 | 22 |
| pneumonia | 57 | 21 | 64 | 107 | 95 | 12 |
| chronic kidney disease (CKD) | 174 | 2 | 8 | 177 | 174 | 3 |
| kidney failure | 102 | 0 | 8 | 108 | 106 | 2 |
| suicide | 0 | 0 | 2 | 2 | 2 | 0 |
| lower respiratory infections (LRI) | 0 | 0 | 42 | 35 | 31 | 4 |
| tuberculosis | 17 | 13 | 29 | 39 | 31 | 8 |

**Supplementary Figures**

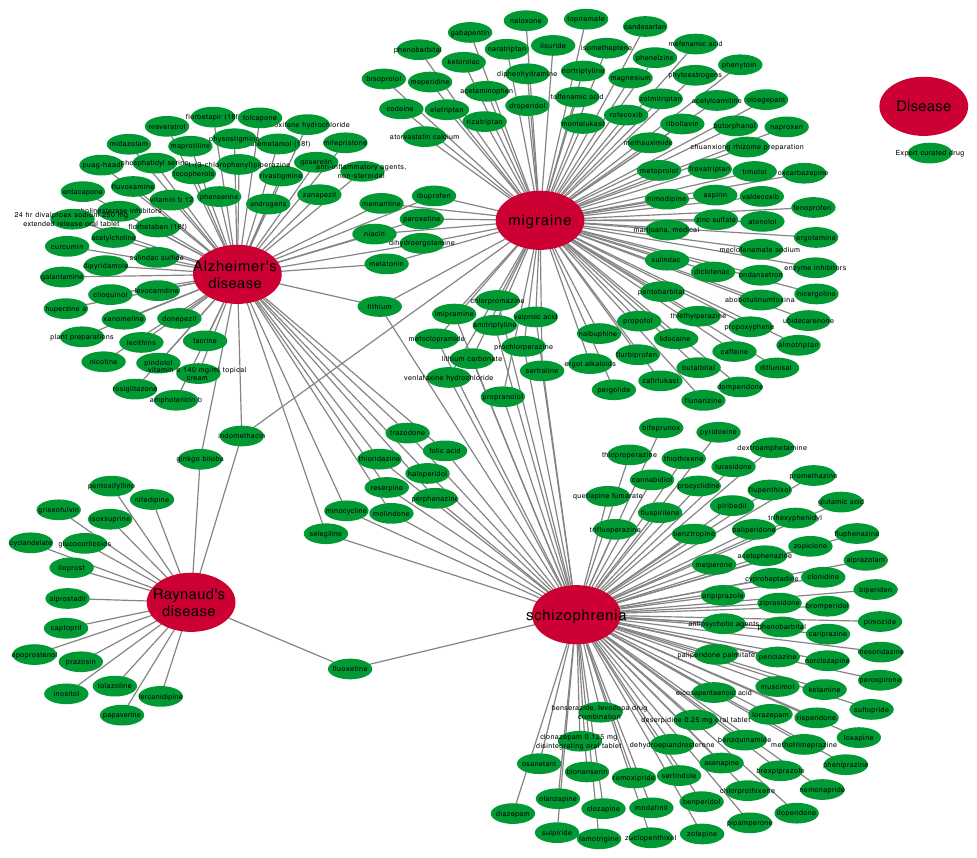

**Supplementary Figure 1.** **Experts curated** **disease-drug associations (DDAs) for RD, migraine, AD, and schizophrenia.** 29 drugs are annotated for two to three diseases.

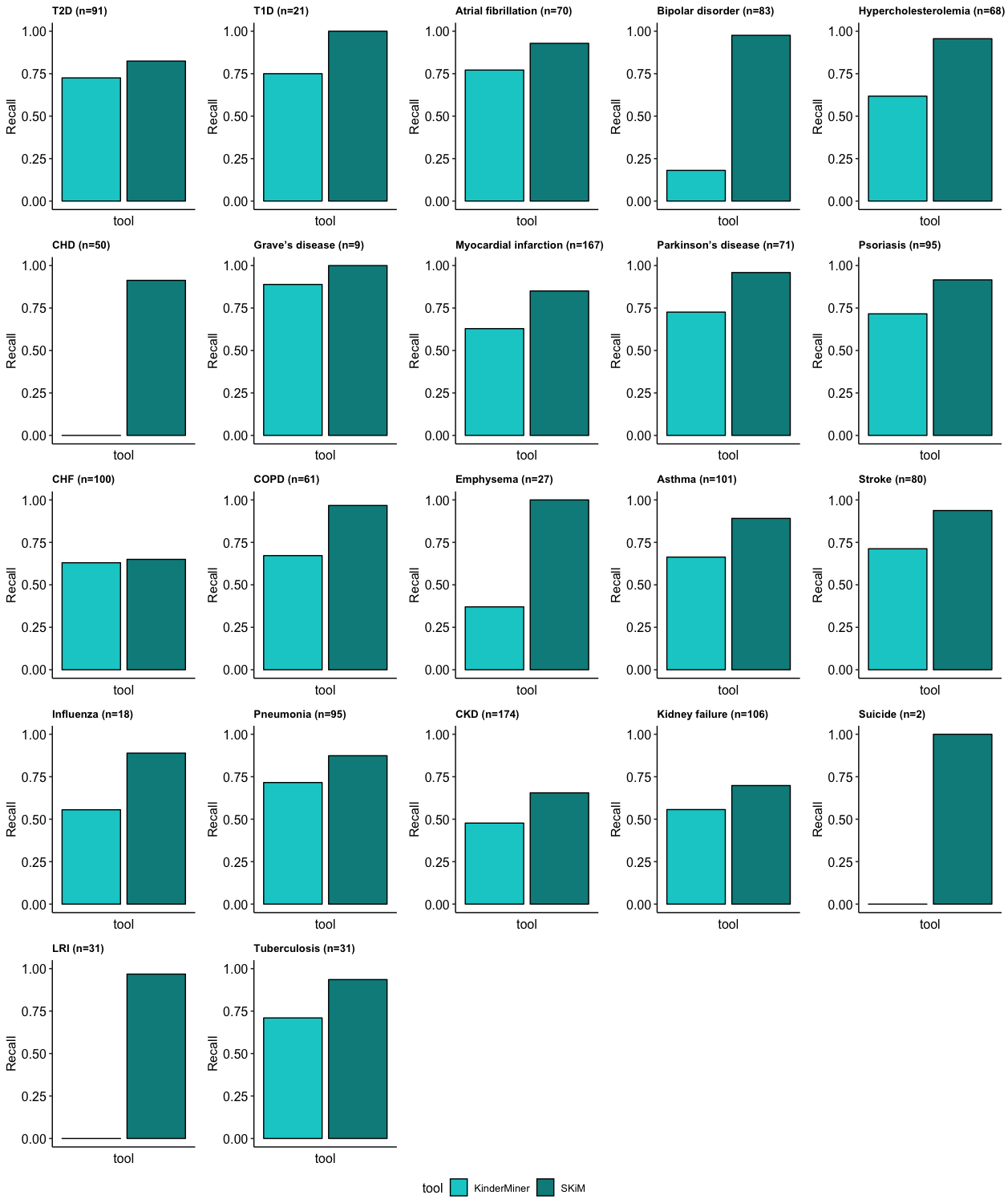

**Supplementary Figure 2. Comparison between recall achieved by SKiM and KinderMiner on experts curated disease-drug associations at cut-off date March 2019.**

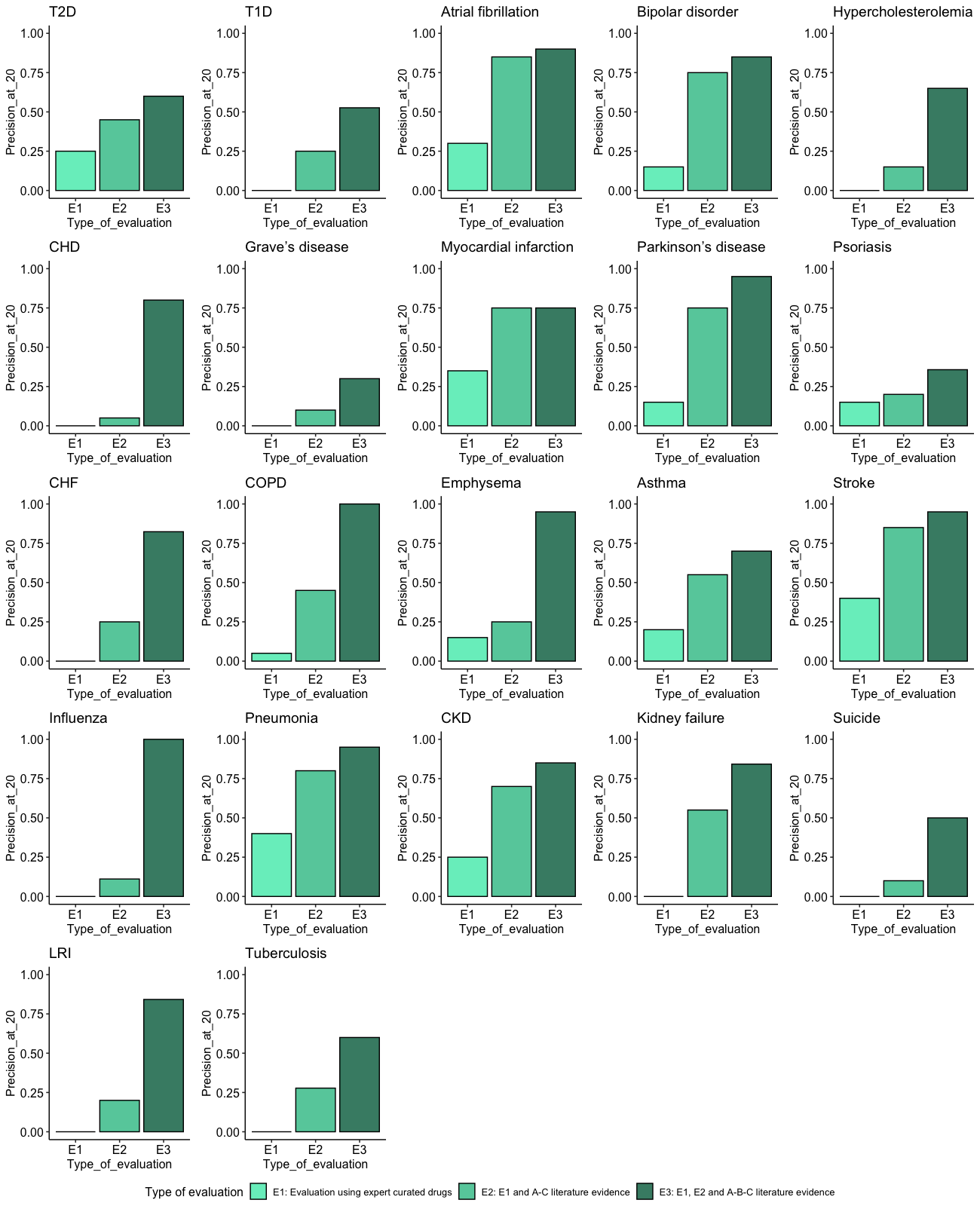

**Supplementary Figure 3. Precision@20 achieved by SKiM for 22 conditions at cut-off date March 2019.**

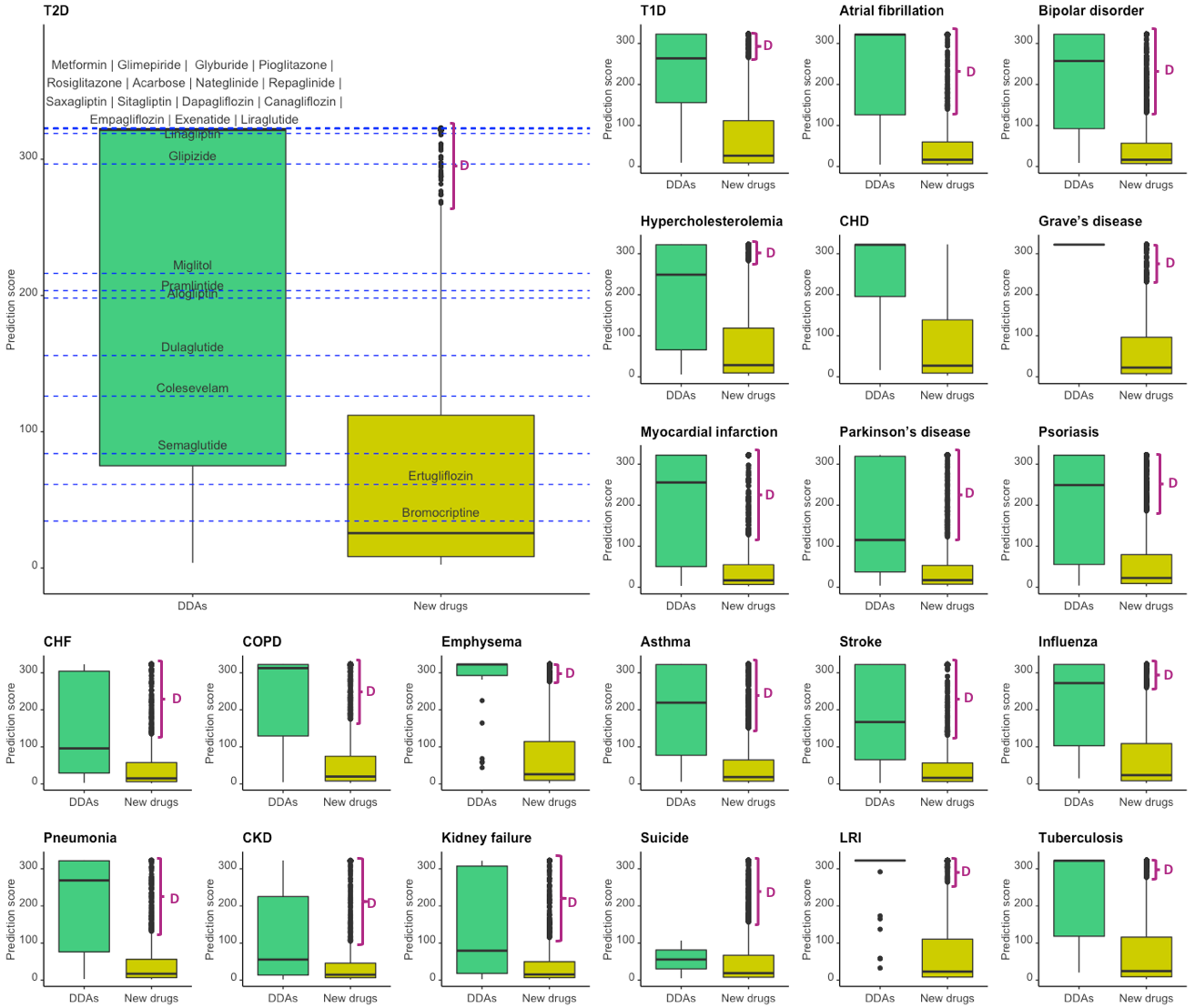

**Supplementary Figure 4. DDAs and new drugs uncovered by SKiM for 22 conditions at cut-off date March 2019.** “D” represents the new drugs with high prediction score. These are promising candidates for wet lab experiments and clinical trials.

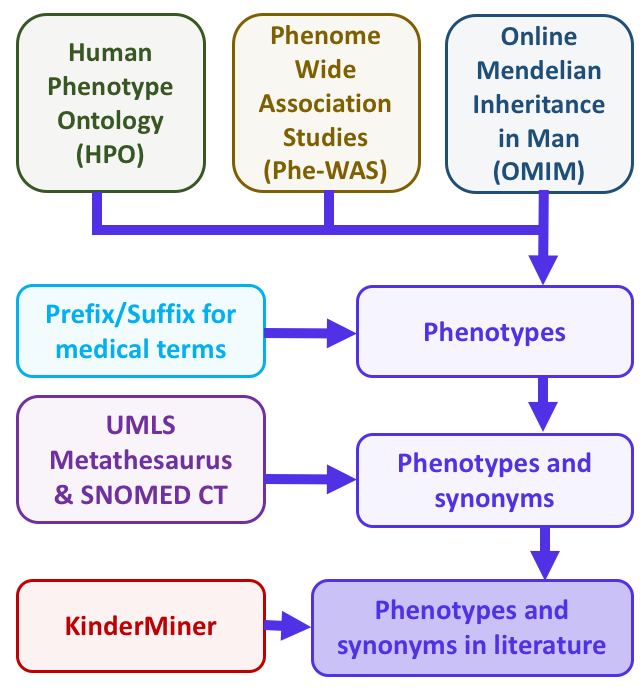

**Supplementary Figure 5. Phenotypes and symptoms lexicon.**

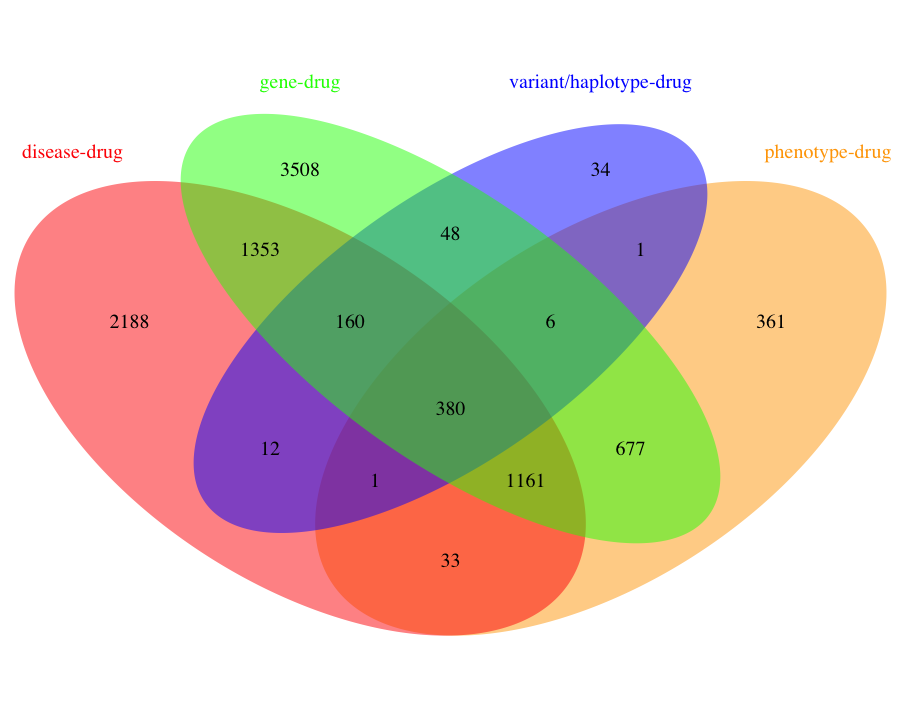

**Supplementary Figure 6.** **Drugs subset from various expert curated associations.**

**Supplementary Methods**

**Supplementary Method 1:** We don't perform ordinary false discovery rate (FDR) correction since the distribution of the FET p-value is discrete and skewed towards 1, violating the model assumptions of most state-of-the-art FDR control methods^18^. We observed that certain terms appear in a small number of PubMed abstracts (e.g. “carbazeran”, a drug used in the treatment of heart failure appears only in 18 abstracts) among ~31M abstracts in PubMed. This violates the model assumptions of most of the state-of-the-art FDR control methods.

We explain the simulation method to find FDR below:

**Simulation settings**

**Marginal settings**

- $N=29613663$ is total number of PMC papers.
- $A$ is disease. $n_{A}$ is the number of occurrence of $A$ in all papers. In the following examples we let $n_{A}=6507$.
- $B_{1},...,B_{9272}$ are symptoms. $C_{1},...,C_{9665}$ are drugs. $n_{B_{i}}$’s and $n_{C_{j}}$’s are their number of occurrence and are stored in separate lists.
- We filter out $B$’s and $C$’s with zero occuurence. There are 6 $B$’s and 65 $C$’s with zero occurrence. After filtering, there are 9266 $B$’s and 9600 $C$’s left.

**Some Notations**

- Between A and B’s: $p_{1}$=P(B occurs | A occurs), $p_{2}$=P(B occurs | A not occurs)
- Between B’s and C’s: $p_{1}$=P(C occurs | B occurs), $p_{2}$=P(C occurs | B not occurs)
- Odds ratio: $OR=\frac{p_{1}/(1-p_{1})}{p_{2}/(1-p_{2})}$

**Define associations (pre-assign true significancy)**

- $A$ is associated with randomly selected 20 of $B_{i}$’s. A is not associated with the rest $B$’s.
- Each $B_{i}$ is associated with randomly selected 10 of $C_{j}$’s. i.e. Among all $9272\times9665$ edges between $B$’s and $C$’s, there are $9272\times10$ significant edges.
- Associated: $OR>=2$ (for simplicity, consider one-sided case)
- Not associated: $OR=1$
- $A$ is associated with $C_{j}$ if $A$ and $C_{j}$ are associated with at least one common $B_{i}$. i.e. we have at most (due to overlapping) $20\times10=200$ $C$’s associated with A among all 9665 $C$’s.

**Model assumptions for simulating contingency table**

For simplicity, take $A\to B_{1}$ as example. All the rest $A\to B_{i}$ and $B_{i}\to C_{j}$ follow the same model with different parameters.

Recall that $p_{1}$=P(B occurs | A occurs), $p_{2}$=P(B occurs | A not occurs). We simulate the number of co-occurrence of $A$ and $B$, $n_{kt}$, from $Binom(p_{1},n_{A})$. We simulate the number of papers where $B$ occurs but $A$ does not occur, $n_{BnotA}$, from $Binom(p_{2},N-n_{A})$.

Note that $n_{kt}+n_{BnotA}$ does not necessarily equal to $n_{B}$, but will be pretty close because the values of $p_{1}$ and $p_{2}$ are calculated based on $n_{B}$ (details in below).

**Parameter values setting**

- Whether associated or not, $p_{2}$ is estimated using the marginal probability. i.e. Between $A$ and $B_{i}$’s: $p_{2}$=P($B_{i}$ occurs)=$n_{B_{i}}/N$. Between $B_{i}$ and $C_{j}$: $p_{2}$=P($C_{j}$ occurs)=$n_{C_{j}}/N$.
- For non-associated pairs: $OR=1$, $p_{1}=p_{2}$.
- For associated pairs: $OR 2+Exp(1)$. $p_{1}$ is calculated using $p_{2}$ and $OR$.
- For associated pairs, in the simulated contingency table, a pseudo-count of min(5, sum of first row, sum of first column) is added to the two slots of co-occurrence and co-unoccurrence and subtracted from the rest two slots . The reason is: for tables with zero co-occurrence, FET tends to overestimate p-value (close to 1) and is underpowered to distinguish them even if they have large OR. For example, if $p_{1}=1e-4$ and $p_{2}=1e-5$, there is a strong association with OR=~10. However, there can easily be tables with zero co-occurrence when $n_{A}$ is not large enough. In our case, $n_{A}=6507$. Expected co-occurrence is $n_{A}\times p_{1}=0.65<1$.
- A pseudo-count could hugely increase $p_{1}$. For example, if $n_{A}=1$, then $p_{1}$ will be 1. This is OK since we only apply to associated pairs. We are just making them more associated so that FET could test them out.

**Our goals**

- Simulate contingency tables with known association and perform FET to get p-values.
- Evaluate the power and FDR under p-value cutoff 1e-5 for
  1. all $A\to B_{i}$ associations
  2. all $B_{i}\to C_{j}$ association for top $N=50$ significant $B_{i}$’s with highest prediction score
  3. all $A\to C_{j}$ associations as defined previously.

**Implementation in R**

**Load data and marginal settings**

```
set.seed(2020)  # for reproducibility
setwd("~/Google Drive/Hallu/codes/ckgroup/SKIM/")
source("simulation_functions.R")

N <- 29613663 # total number of papers in database
n_A <- 6507 # number of papers containing A
meta_B <- readxl::read_xlsx("SKiM_Files/To_Zijian/Phenotypes_and_symptoms_count.xlsx", sheet = 1)
meta_C <- readxl::read_xlsx("SKiM_Files/To_Zijian/Drugs_count.xlsx",sheet = 1)
n_B <- meta_B$Phenotype_and_symptom_count # number of papers containing each B
n_C <- meta_C$Drug_count # number of papers containing each C

# Filter out zero occurrence
meta_B <- meta_B[n_B>0,]
n_B <- n_B[n_B>0]
meta_C <- meta_C[n_C>0,]
n_C <- n_C[n_C>0]

p_B <- n_B/N
p_C <- n_C/N
```

Check the metadata, counts and probability of $B$’s:

```
head(meta_B)
## # A tibble: 6 x 3
##   Phenotype_and_symptom                 Phenotype_and_symptom_~ db_article_count
##   <chr>                                                   <dbl>            <dbl>
## 1 C0239337_T190:abnormal_limbs                             1441         29613663
## 2 M00004289_T033:abnormality_of_the_po~                     100         29613663
## 3 C0596875_T047:lysinemia                                     2         29613663
## 4 C1843112_T033:broad_nail                                    2         29613663
## 5 D00000686_T777:normal_cry                                   3         29613663
## 6 C1859698_T033:large_joint_contractur~                      39         29613663
head(n_B)
## [1] 1441  100    2    2    3   39
head(p_B)
## [1] 4.865997e-05 3.376820e-06 6.753639e-08 6.753639e-08 1.013046e-07
## [6] 1.316960e-06
```

**Define associations**

```
#######################################
# Set significant B's
n_signif_B <- 20 # number of significant B's

# indices of significant B's to A
which_signif_BtoA <- sample.int(nrow(meta_B), n_signif_B)

#######################################
# Set significant C's for each B
n_signif_C <- 10 # number of significant C's for each B
# indices of significant C's to each B, as columns
which_signif_CtoB <- replicate(nrow(meta_B),sample.int(nrow(meta_C), n_signif_C))

#######################################
# Set significant C's to A
which_signif_CtoA <- unique(as.vector(which_signif_CtoB[,which_signif_BtoA]))
```

Overview of assigned associated terms:

```
str(which_signif_BtoA)
##  int [1:20] 7767 8920 4417 8465 170 7878 945 4992 2602 3062 ...
str(which_signif_CtoB)
##  int [1:10, 1:9266] 1524 7468 8827 8182 4690 3142 5163 2509 1404 4952 ...
str(which_signif_CtoA)
##  int [1:198] 8280 8779 8358 1978 946 4271 7793 2368 4291 9159 ...
```

**Parameter setting for** $\boldsymbol{A\to}\boldsymbol{B}_{\boldsymbol{i}}$**’s**

```
#######################################
# Set p_1 and p_2 for A and each B
p_1_BtoA <- p_2_BtoA <- p_B
OR_BtoA <- 2+rexp(n_signif_B)
p_1_BtoA[which_signif_BtoA] <- get_p1(p_2_BtoA[which_signif_BtoA],OR_BtoA)
```

Check $p_{1}$ and $p_{2}$ for associated $A\to B_{i}$’s:

```
p_1_BtoA[which_signif_BtoA]
##  [1] 3.349202e-06 2.540042e-07 1.466906e-06 3.438339e-06 8.673815e-04
##  [6] 4.466043e-06 1.366215e-04 1.635897e-06 3.494111e-05 1.037920e-05
## [11] 6.905228e-04 1.797883e-06 1.238637e-05 1.169223e-03 9.842889e-06
## [16] 6.070601e-05 1.631564e-05 1.527495e-05 3.804508e-06 2.169919e-05
p_2_BtoA[which_signif_BtoA]
##  [1] 9.455095e-07 1.013046e-07 6.753639e-07 1.215655e-06 3.692215e-04
##  [6] 1.587105e-06 2.232078e-05 4.389866e-07 1.266307e-05 3.376820e-06
## [11] 2.922975e-04 6.078275e-07 6.179580e-06 3.921501e-04 1.553337e-06
## [16] 2.775746e-05 7.395235e-06 7.125090e-06 1.891019e-06 6.280885e-06
```

**Simulate contingency table and perform FET for all** $\boldsymbol{A\to}\boldsymbol{B}_{\boldsymbol{i}}$**’s**

```
#######################################
# Simulate contingency tables, calculate sort ratios and perform FET between A and each B
pval_BtoA <- sort_ratio_BtoA <- numeric(nrow(meta_B))

for(B_idx in seq_along(pval_BtoA)){
    temp_table <- simulate_table(N, n_A, N-n_A, p_1_BtoA[B_idx], p_2_BtoA[B_idx])
    # Add pseudo-count for true associations
    if(B_idx%in%which_signif_BtoA){
      pseudo_mat <- min(5,sum(temp_table[,1]),sum(temp_table[1,]))*
        matrix(c(1,-1,-1,1),2,2)
      temp_table <- temp_table+pseudo_mat
    }
    sort_ratio_BtoA[B_idx] <- temp_table[1,1]/sum(temp_table[,1])
    pval_BtoA[B_idx] <- fisher.test(temp_table)$p.value
}

# Calculate prediction score
score_BtoA <- -log10(pval_BtoA)+log10(sort_ratio_BtoA)
```

Example table with association:

```
simulate_table(N, n_A, N-n_A,
               p_1_BtoA[which_signif_BtoA[1]],
               p_2_BtoA[which_signif_BtoA[1]])+
  5*matrix(c(1,-1,-1,1),2,2)
##        have_y     no_y
## have_x      5     6502
## no_x       19 29607137
```

Example table without association:

```
simulate_table(N, n_A, N-n_A, p_1_BtoA[1], p_2_BtoA[1])
##        have_y     no_y
## have_x      0     6507
## no_x     1397 29605759
```

Check p-value of associated ones and randomly selected 10:

```
pval_BtoA[which_signif_BtoA]
##  [1] 1.024798e-13 1.060390e-11 2.707238e-14 5.005108e-14 4.485487e-05
##  [6] 8.688404e-13 1.144461e-08 2.229382e-15 2.013313e-08 2.914793e-11
## [11] 8.105453e-04 3.157713e-15 4.478054e-10 3.290097e-04 2.320763e-12
## [16] 1.435043e-06 1.197714e-11 2.017229e-09 5.514549e-13 9.174042e-10
sample(pval_BtoA,10)
##  [1] 0.1109063 1.0000000 1.0000000 1.0000000 1.0000000 1.0000000 1.0000000
##  [8] 1.0000000 1.0000000 0.7860476
```

Number of p-values less or equal to 1e-5:

```
sum(pval_BtoA<=1e-5)
## [1] 17
```

**Select top** $\boldsymbol{N=50}$ $\boldsymbol{B}$**’s with highest prediction score among significant** $\boldsymbol{B}$**’s**

Since we have less than $N$ significant $B$’s, we just use these significant ones instead of 50.

```
######################################
# Keep significant Bs with p-value <=1e-5, then
# choose top 50 B's with largest prediction score
# or largest p-values

#cand_B <- which(rank(pval_BtoA)<=50)
cand_B <- which(rank(-score_BtoA)<=50)

# Since there are only 20 pval left, just use these 20 instead of 50.
cand_B <- cand_B[pval_BtoA[cand_B]<=1e-5]
```

Indices of candidate $B$’s to keep:

```
cand_B
##  [1]   78  724  945 2602 3062 3662 4417 4483 4684 4992 5104 6888 7687 7767 7878
## [16] 8465 8920
```

For stress test, we can also use top $N=50$ $B$’s with smallest p-values regardless of their significancy.

cand_B_2 <- **which**(**rank**(pval_BtoA)**<=**50)

**Simulate contingency table and perform FET for top** $\boldsymbol{N}$ $\boldsymbol{B}$**’s and all** $\boldsymbol{C}$**’s**

For significant $B$’s:

res_CtoB <- **test_CtoB**(cand_B,p_C,which_signif_CtoB, verbose=F)

A matrix of p-values:

```
str(res_CtoB$PVAL_CtoB)
##  num [1:9600, 1:17] 1 1 1 1 1 1 1 1 1 1 ...
##  - attr(*, "dimnames")=List of 2
##   ..$ : NULL
##   ..$ : chr [1:17] "78" "724" "945" "2602" ...
```

Number of significant $C$’s for $B_{1}$:

```
sum(res_CtoB$PVAL_CtoB[,1]<=1e-5)
## [1] 9
```

For top 50 $B$’s:

```
##  num [1:9600, 1:50] 1 1 1 1 1 1 1 1 1 1 ...
##  - attr(*, "dimnames")=List of 2
##   ..$ : NULL
##   ..$ : chr [1:50] "78" "170" "303" "724" ...
```

**Evaluate power and FDR by comparing predicted association (p-value<=1e-5) with true association**

**Evaluate for predicted significancy between all** $\boldsymbol{A\to}\boldsymbol{B}_{\boldsymbol{i}}$**’s**

```
get_power_FDR(pval_BtoA<=1e-5, seq_len(nrow(meta_B))%in%which_signif_BtoA)
## $power
## [1] 0.85
##
## $FDR
## [1] 0
```

**Evaluate for predicted significancy between top** $\boldsymbol{N}$ $\boldsymbol{B}$**’s and all** $\boldsymbol{C}$**’s**

For significant $B$’s:

```
# True significancy
true_signif_CtoB <- apply(which_signif_CtoB[,cand_B],
                          2, function(x) seq_len(nrow(meta_C))%in%x)

get_power_FDR(as.vector(res_CtoB$PVAL_CtoB<=1e-5), as.vector(true_signif_CtoB))
## $power
## [1] 0.9823529
##
## $FDR
## [1] 0
```

For top 50 $B$’s:

```
# True significancy
true_signif_CtoB_2 <- apply(which_signif_CtoB[,cand_B_2],
                          2, function(x) seq_len(nrow(meta_C))%in%x)

get_power_FDR(as.vector(res2_CtoB$PVAL_CtoB<=1e-5), as.vector(true_signif_CtoB_2))
## $power
## [1] 0.93
##
## $FDR
## [1] 0.002145923
```

**Evaluate for predicted significancy between** $\boldsymbol{A}$ **and** $\boldsymbol{C}$**’s**

For significant $B$’s:

```
SKiM_signif_CtoA <- apply(res_CtoB$PVAL_CtoB,1,function(x) any(x<=1e-5))

get_power_FDR(SKiM_signif_CtoA, seq_len(nrow(meta_C))%in%which_signif_CtoA)
## $power
## [1] 0.8333333
##
## $FDR
## [1] 0
```

For top 50 $B$’s (same result since only significant $B$’s will be used to link between $A$ and $C$’s):

```
SKiM_signif_CtoA_2 <- apply(res_CtoB$PVAL_CtoB,1,function(x) any(x<=1e-5))

get_power_FDR(SKiM_signif_CtoA_2, seq_len(nrow(meta_C))%in%which_signif_CtoA)
## $power
## [1] 0.8333333
##
## $FDR
## [1] 0
```

**Why FDR is zero**

- The reason of zero FDR is under such biased contigency tables, the p-value distribution of FET under null hypothesis (no association) is discrete and highly skewed towards 1. It’s hard to observe small p-values for un-associated pairs. See discussions in [this paper](https://www.microsoft.com/en-us/research/wp-content/uploads/2016/02/fdr20for20contingency20tables.pdf). When the p-value cutoff is conservative enough (1e-5 in SKiM), we are not making any false positives.

**More evaluations with random replications**

We now test on a wide range of common and rare diseases. Candidate values of $n_{A}$ are 500, 5,000, 50,000, 100,000, 200,000. For each $n_{A}$, we repeat 10 times of the simulation and report the average power and FDR. Codes for this part are stored in a separate file. We just show the final results here:

```
sim_out <- read.csv("sim_out_10012020.csv")
sim_out
##   X       n_A power_AtoB power_BtoC power_AtoC FDR_AtoB    FDR_BtoC    FDR_AtoC
## 1 1   n_A=500       0.96  0.9395182  0.9026694        0 0.002295158 0.002323967
## 2 2  n_A=5000       0.90  0.9562515  0.8619289        0 0.001618715 0.001651652
## 3 3 n_A=50000       0.83  0.9504965  0.7887817        0 0.000000000 0.000000000
## 4 4 n_A=1e+05       0.76  0.9403226  0.7157341        0 0.002638067 0.002646379
## 5 5 n_A=2e+05       0.77  0.9272452  0.7169642        0 0.003433889 0.003457114
```

**Session Information**

```
sessionInfo()
## R version 4.0.2 (2020-06-22)
## Platform: x86_64-w64-mingw32/x64 (64-bit)
## Running under: Windows 10 x64 (build 18363)
##
## Matrix products: default
##
## locale:
## [1] LC_COLLATE=Chinese (Simplified)_China.936
## [2] LC_CTYPE=Chinese (Simplified)_China.936
## [3] LC_MONETARY=Chinese (Simplified)_China.936
## [4] LC_NUMERIC=C
## [5] LC_TIME=Chinese (Simplified)_China.936
##
## attached base packages:
## [1] stats     graphics  grDevices utils     datasets  methods   base
##
## other attached packages:
## [1] qvalue_2.21.0
##
## loaded via a namespace (and not attached):
##  [1] Rcpp_1.0.5       pillar_1.4.6     compiler_4.0.2   cellranger_1.1.0
##  [5] plyr_1.8.6       tools_4.0.2      digest_0.6.25    evaluate_0.14
##  [9] lifecycle_0.2.0  tibble_3.0.3     gtable_0.3.0     pkgconfig_2.0.3
## [13] rlang_0.4.7      cli_2.0.2        yaml_2.2.1       xfun_0.18
## [17] stringr_1.4.0    dplyr_1.0.2      knitr_1.30       generics_0.0.2
## [21] vctrs_0.3.4      grid_4.0.2       tidyselect_1.1.0 glue_1.4.2
## [25] R6_2.4.1         fansi_0.4.1      readxl_1.3.1     rmarkdown_2.4
## [29] ggplot2_3.3.2    purrr_0.3.4      reshape2_1.4.4   magrittr_1.5
## [33] scales_1.1.1     ellipsis_0.3.1   htmltools_0.5.0  splines_4.0.2
## [37] assertthat_0.2.1 colorspace_1.4-1 utf8_1.1.4       stringi_1.5.3
## [41] munsell_0.5.0    crayon_1.3.4
```

**Supplementary Method 2:** Automated time-slicing approach: Evaluating SKiM only on the discoveries by Swanson and his colleagues is not sufficient. LBD systems uncover new discoveries that are yet to be validated, and evaluating their performance is challenging^19^. A recent work suggests an automated time-slicing approach for evaluating LBD systems like SKiM^20^. In this approach, PubMed is divided into two segments, pre-cut-off and post-cut-off at a cut-off date. A→C from both the segments is retrieved, and Cs present only in the post-cut-off are considered as the gold standard. The performance of LBD systems is evaluated on the gold standard at the same cut-off date and reported with the standard evaluation metrics, recall, and precision. We used the automated time-slicing approach to generate gold standards for four diseases from the discoveries of Swanson and his colleagues: RD, migraine, AD, and schizophrenia (Supplementary Method Table 1).

**Supplementary Method Table 1.** Gold standard

| **Disease** | **Cut-off date** | **Drugs in**  **pre-cut-off (I_pre_)** | **Drugs in**  **post-cut-off (I_post_)** | **G = I_post_ - I_pre_** |
| --- | --- | --- | --- | --- |
| RD | 1985 | 37 | 96 | **70** |
| migraine | 1987 | 64 | 195 | **144** |
| AD | 1995 | 285 | 442 | **292** |
| schizophrenia | 1997 | 121 | 249 | **158** |

Note: G – Gold standard drugs count.

We evaluated SKiM findings using the gold standard. SKiM achieved a recall between 0.5685 to 0.70 at FET p-value less than 1.0×10^-05^ (see R in Supplementary Method Table 2). We observed that five drugs for RD, 19 drugs for migraine, 30 drugs for AD, and 20 drugs for schizophrenia are mentioned for the first time in PubMed abstracts only after the cut-off date. Thus, it is impossible to predict these drugs using PubMed articles published up to the cut-off date. We recalculated false negatives (see FN1 in Supplementary Method Table 2) by excluding the drugs mentioned in PubMed abstracts only after the cut-off date. SKiM achieved a recall between 0.6336 to 0.7539 at FET p-value less than 1.0×10^-05^ (see R1 in Supplementary Method Table 2).

**Supplementary Method Table 2.** Performance of SKiM at FET p-value less than 1.0×10^-05^

| **Disease** | **Cut-off date** | **G** | **SKiM** | | | | |
| --- | --- | --- | --- | --- | --- | --- | --- |
|  |  |  | **TP** | **FN** | **R** | **FN1** | **R1** |
| RD | 1985 | 70 | 49 | 21 | **0.70** | 16 | **0.7539** |
| migraine | 1987 | 144 | 88 | 56 | **0.6111** | 37 | **0.704** |
| AD | 1995 | 292 | 166 | 126 | **0.5685** | 96 | **0.6336** |
| schizophrenia | 1997 | 158 | 98 | 60 | **0.6203** | 40 | **0.7101** |

Notes: G – Gold standard drugs count; TP – True positive; FN – False negative; R – Recall from TP and FN.

FN1 – Five drugs for RD, 19 drugs for migraine, 30 drugs for AD, and 20 drugs for schizophrenia are mentioned in PubMed after the cut-off date. These drugs were excluded from FN to give FN1; R1 – Recall from TP and FN1.

We relaxed FET p-value cut-off to 0.05 to see the performance of SKiM findings against the gold standard. True positives (see TP in Supplementary Method Table 3) increased and false negatives (see FN in Supplementary Method Table 3) decreased when compared to true positives (see TP in Supplementary Method Table 2) and false negatives (see FN in Supplementary Method Table 2) obtained at FET p-value less than 1.0×10^-05^ (see TP and FN in Supplementary Method Table 2). SKiM achieved a recall between 0.7153 to 0.7774 at FET p-value less than 0.05 (see R in Supplementary Method Table 3). Similar to the performance at FET p-value less than 1.0×10^-05^, we excluded five drugs for RD, 19 drugs for migraine, 30 drugs for AD, and 20 drugs for schizophrenia mentioned in PubMed abstract only after the cut-off date and recalculated false negatives (see F2 in Supplementary Method Table 3). SKiM achieved a recall between 0.8 to 0.8664 at FET p-value less than 0.05 (see R2 in Supplementary Method Table 3). False negatives (see Supplementary Method Table 1, Supplementary Method Table 2, and Supplementary Method Table 3) are due to weak or no association of A or C terms with the intermediate B terms. The precision is expected to be low because A🡪B🡪C retrieves all possible Cs associated with multiple Bs. On the other hand, the gold standard includes Cs that are associated with only with A (i.e. A🡪C association).

**Supplementary Method Table 3.** Performance of SKiM at FET p-value less than 0.05

| **Disease** | **Cut-off date** | **G** | **SKiM** | | | | |
| --- | --- | --- | --- | --- | --- | --- | --- |
|  |  |  | **TP** | **FN** | **R** | **FN2** | **R2** |
| RD | 1985 | 70 | 52 | 18 | **0.7429** | 13 | **0.8** |
| migraine | 1987 | 144 | 103 | 41 | **0.7153** | 22 | **0.824** |
| AD | 1995 | 292 | 227 | 65 | **0.7774** | 35 | **0.8664** |
| schizophrenia | 1997 | 158 | 118 | 40 | **0.7468** | 20 | **0.8551** |

Notes: G – Gold standard drugs count; TP – True positive; FN – False negative; R – Recall from TP and FN.

FN2 – Five drugs for RD, 19 drugs for migraine, 30 drugs for AD, and 20 drugs for schizophrenia are mentioned in PubMed after the cut-off date. These drugs were excluded from FN to give FN2; R2 – Recall from TP and FN2.

Automated time-slicing approach is for generating a gold standard for evaluating LBD systems. The approach has been applied to compare various ranking algorithms based on association rules, term frequency-inverse document frequency, Z-score, or mutual information measure using an existing system LitLinker^20,21^. To our knowledge, none of the existing LBD systems including LitLinker were evaluated on automated time-slicing approach. Most of the existing LBD systems are not functional and cannot be evaluated on the gold standard generated by us using the automated time-slicing approach. Though BITOLA^22^ and LION LBD^4^ are functional, the systems are not reliable for drug repurposing (see Introduction section for details). Thus, a comparison between SKiM and the existing LBD systems on an automated time-slicing approach is not possible.

**Supplementary Discussion**

**Supplementary Discussion 1:** In the current study, we took top n=50 phenotypes and symptoms from A🡪B for executing B🡪C. Though our approach of selecting top n B terms is based on existing LBD systems, the value of n is arbitrary and the number of C terms uncovered by SKiM depends on n. SKiM is expected to give additional C terms when all Bs from A🡪B are considered for executing B🡪C. However, all phenotypes and symptoms from A🡪B may not be associated with drugs. This is true for the execution with top 50 phenotypes and symptoms from A🡪B (e.g. finger swelling and acrocyanosis for RD, and transient global amnesia and scintillating scotoma for migraine).
